# Retrieval practice is associated with attenuated stress-related inference impairment and stronger reactivation of bridge-related category information

**DOI:** 10.1101/2025.03.14.643187

**Authors:** Jinpeng Guo, Ruixin Chen, Qi Zhao, Xiaojun Sun, Wei Liu

## Abstract

A hallmark of human memory is its ability to form novel inferences by linking discrete but related events. We examined whether acute stress impairs memory inference and if retrieval practice can buffer this effect. Participants were trained on image pairs, AB and BC, to establish interconnected triads (ABC) with a shared bridge element B. Twenty-four hours later, we induced acute stress in half of the participants and then tested their capacity to infer the indirect AC associations. Behavioral results indicated that acute stress reduced inference accuracy and slowed responses, and retrieval practice was associated with attenuation of stress-induced impairment in A-C inference accuracy and speed. Using multivariate decoding analysis of human electroencephalogram (EEG) recordings, we found neural evidence of rapid reactivation of bridge-related category information during inference, a neural signature predictive of subsequent successful inference. Importantly, stress was associated with reduced bridge-related category reactivation, whereas retrieval practice was associated with stronger reactivation under stress. Collectively, our findings link rapid reactivation of bridge-related category information to successful memory-based inference. Although susceptible to stress, this reactivation was stronger following retrieval practice, suggesting that robust memory traces may contribute to more resilient inference under stress.

**Significance Statement:** Stress often disrupts our ability to form new connections from past experiences, thereby limiting cognitive flexibility. Our study shows that stress impairs memory inference—the ability to link related events to generate novel links—and is associated with reduced rapid reactivation of bridge-related category information. However, we found that a simple yet effective technique, retrieval practice, which involves recalling related information, is associated with mitigating this negative effect. Reactivating prior memory traces through repeated recall was associated with reduced inference impairment and even stronger memory reactivation signals. These findings highlight the potential of using strategic learning methods to protect and enhance cognitive flexibility, particularly under stress, with broad implications for improving memory resilience in high-pressure environments such as examinations.

## Introduction

The ability to infer novel relationships from distinct yet related memories is essential for adaptive behavior. This capacity allows individuals to integrate separate experiences—such as linking a new event with a stored memory—to form novel indirect associations. Theoretical models suggest that successful inference relies on a specific constructive mechanism during retrieval: the rapid reactivation of the shared “bridge” element that links discrete episodes. However, acute stress is known to impair memory retrieval, potentially disrupting this delicate inferential process (Vogel and Schwabe, 2016). Conversely, retrieval practice is well-established to strengthen memory traces and facilitate knowledge transfer, potentially offering a countermeasure to stress-related deficits. In the present study, we examined how acute stress impairs the cognitive and neural mechanisms underlying memory inference. Furthermore, we examined whether retrieval practice was associated with reduced stress-related inference impairment and bridge-related reactivation across experiments.

To investigate the neural underpinnings of memory inference, we recorded electroencephalogram (EEG) signals while participants performed an associative inference task (Preston and Eichenbaum, 2013; Schlichting and Preston, 2015). This paradigm requires recombining information across overlapping pairs (e.g., AB, BC) to make a novel inference (e.g., A-C). Previous fMRI studies have highlighted the roles of the prefrontal cortex (PFC) and medial temporal lobe (MTL) in integrating these overlapping associations (Zeithamova and Preston, 2010; Zeithamova et al., 2012). More recently, multivariate decoding analyses have begun to elucidate how the representation of the linking “bridge element” (i.e., B) is reactivated to support subsequent inference (Morton et al., 2020; Schlichting et al., 2021). However, because these studies often lacked neural data during the active A-C inference test, the direct role of the bridge element in supporting online inference remains unclear. We posited that the rapid neural reactivation of the bridge element B is crucial for successful inference. By leveraging the high temporal resolution of EEG and employing time-resolved multivariate decoding analysis (Cichy et al., 2014; Linde-Domingo et al., 2019), we aimed to capture this reactivation during the inferential process itself, focusing on distinct stimulus categories (i.e., faces and buildings) as bridge elements.

Acute stress is known to disrupt episodic memory retrieval, often by affecting the prefrontal-hippocampal circuit (de Quervain et al., 1998; Buchanan et al., 2006; Wolf, 2017; Gagnon et al., 2019). Importantly, under stress, individuals tend to shift from flexible, MTL-dependent cognitive strategies to more rigid, striatal-dependent habitual behaviors (Schwabe et al., 2022), such as relying on familiar paths rather than discovering novel shortcuts during navigation tasks (Brown et al., 2020). Given these neurocognitive alterations, we anticipated that acute stress would hinder the flexible recombination of stored memories (i.e., AB and BC) required for inference (i.e., from A to C).

Considering the detrimental effect of stress on memory, we sought to identify a strategy to protect inferential abilities. While various methods have been explored to mitigate stress-induced memory deficits (J-F de Quervain et al., 2007; Schwabe et al., 2009; Smith and Thomas, 2018), we focused on retrieval practice—the act of actively testing memory. Retrieval practice is a highly effective learning technique that not only enhances overall memory (Karpicke and Blunt, 2011; Roediger and Butler, 2011) but has also been shown to prevent stress-induced impairments in free recall (Smith et al., 2016). This protective effect may arise because retrieval practice promotes rapid consolidation (Antony et al., 2017; Liu et al., 2020; Zhuang et al., 2021), creating stress-resilient cortical memory representations that are less susceptible to stress-induced glucocorticoid release (De Quervain et al., 2003).

The present study integrates these theoretical frameworks. To isolate stress effects specifically on memory retrieval (Kuhlmann et al., 2005; Schönfeld et al., 2014; Smith et al., 2016) and allow the robust long-term benefits of retrieval practice to emerge (Roediger and Karpicke, 2006), we adopted a standard 24-hour delay paradigm (learning and practice on Day 1; stress induction and inference testing on Day 2). We used EEG decoding, along with behavioral measures of inference accuracy and speed in humans, to test a series of hypotheses (**Figure 1**). First, we hypothesized that the rapid neural reactivation of the bridge element B supports successful memory inference. Second, we predicted that acute stress would disrupt this neural reactivation process and impair inferential performance. Finally, we separated our retrieval-practice prediction into two hypotheses. Behaviorally, we hypothesized that retrieval practice of the directly learned AB and BC premise pairs would attenuate stress-related impairment in later A-C inference. Neurally, we hypothesized that retrieval practice would be associated with preservation of bridge-related memory reactivation under stress.

**Figure 1.**
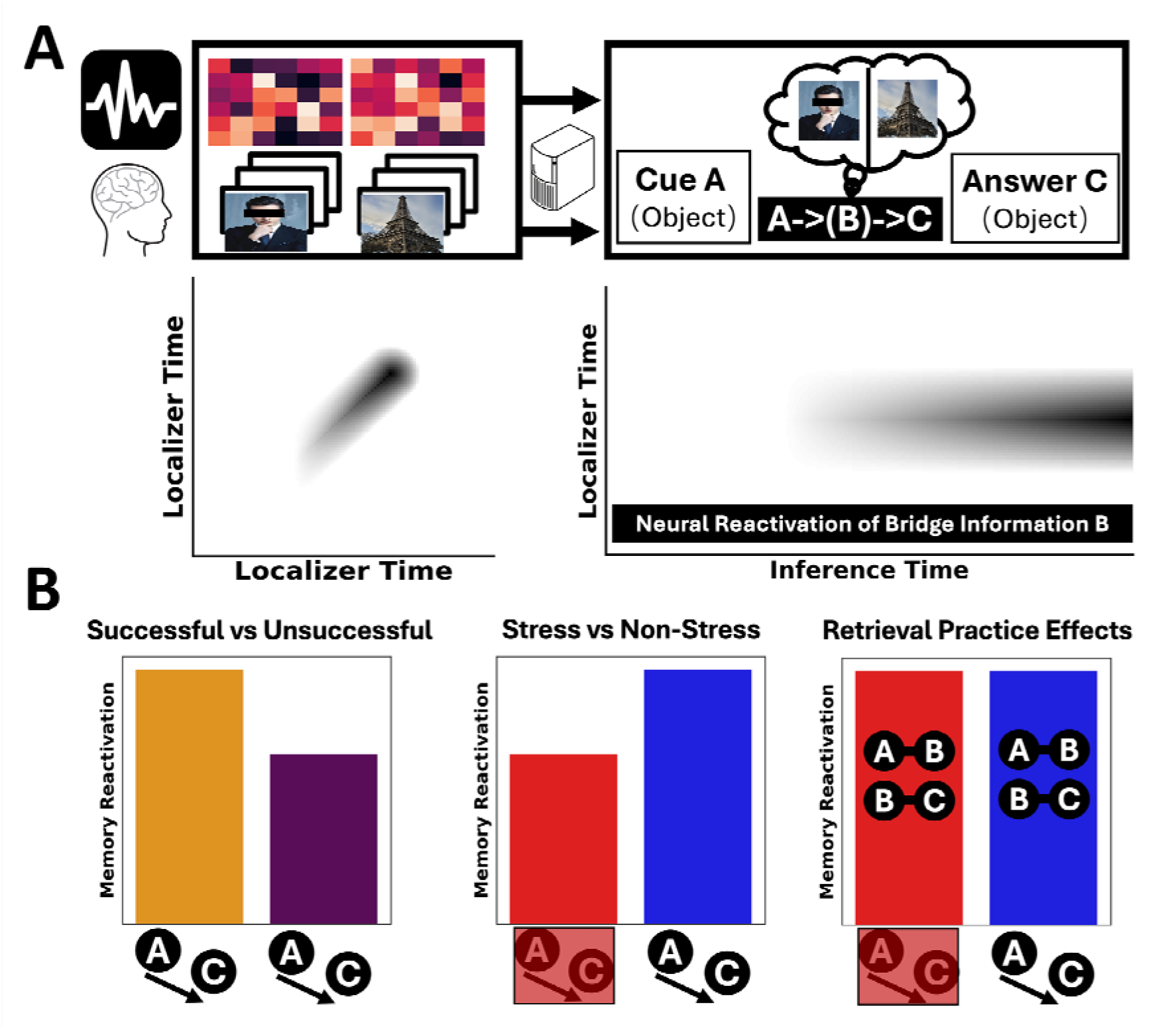
Design for EEG experiment and neural decoding for memory reactivation. **(A)** In the EEG experiment, participants performed an inference task in which they were asked to infer the correct answer-C based on the presented cue-A, using pretrained AB and BC associations. Element B, the “bridge information,” could be either a face or a building image. Both the cue-A and the answer-C were pictures of daily-life objects. EEG data were recorded during both the inference and an independent localizer phase. During the localizer phase, participants actively viewed a different set of face and building images. A classifier was trained on category-specific neural response patterns from the localizer phase and then applied to the inference phase. This approach enabled time-series decoding analyses to detect, within individual trials, whether category information of the bridge element-B was reactivated at any moment. **(B)** Schematic illustration of the main predictions for bridge-related memory reactivation during A-C inference. The y-axis in each panel represents predicted reactivation strength. Left: bridge-related reactivation was expected to be stronger before successful than unsuccessful A-C inference responses (orange = successful trials; purple = unsuccessful trials). Middle: acute stress was expected to reduce bridge-related reactivation relative to the non-stress control condition (red = stress; blue = non-stress control). Right: retrieval practice of the directly learned AB and BC premise pairs was expected to attenuate the stress-related reduction in bridge-related reactivation. The icons below each panel indicate the corresponding A-C inference outcome or experimental group comparison.

## Method

### 2.1 Participants

We tested a total of 136 healthy college students across both experiments. Experiment 1 involved 68 participants (50 females; M_age_ = 20.43 ± 1.80 years), who were randomly assigned to either a stress condition (n = 34; M_age_ = 20.35 years, SD_age_ = 1.87, 26 females) or a control condition (n = 34; M_age_ = 20.50 years, SD_age_ = 1.75, 24 females). There were no significant differences in age or gender between the two conditions (age: t(66) = −0.33, *p* = 0.74, Cohen’s d = −0.08, 95% CI = [−0.56, 0.39]; gender: χ^2^(1,68) = 0.08, *p* = 0.78). The remaining 68 participants (59 females; M_age_ 20.16 ± 1.53 years) were tested in the Experiment 2, which involved random assignment to inference under stress (n = 34; M_age_ = 20.03 years, SD_age_ = 1.51, 32 females) or inference without stress (n = 34; M_age_ = 20.29 years, SD_age_ = 1.57, 27 females). Again, there were no significant differences in age or gender between these groups (age: t(66) = −0.71, *p* = 0.48, Cohen’s d = −0.17, 95% CI = [−0.65, 0.30]; gender: χ^2^(1,68) = 2.05, *p*=0.15). All participants were right-handed and had normal or corrected-to-normal vision. We selected our group sizes based on established precedent in studies using the AB–BC memory inference paradigm (Backus et al., 2016). These sample sizes also fall within the range typically adopted in between-group studies examining stress-related effects on memory (Qin et al., 2009; van Marle et al., 2009; Smith et al., 2016; Vogel et al., 2018). A formal a priori power analysis was not conducted, as the effect size for the interaction between acute stress and retrieval practice on neural markers of inference was not known. Our sample size is comparable to or larger than those used in prior related studies. A post-hoc sensitivity analysis confirmed that our sample was sufficiently large to detect the critical stress-induced impairment on inference performance (observed effect size Cohen’s d = 0.56). The study received approval from the Ethics Review Committee for Life Sciences at Central China Normal University, and all participants provided informed consent prior to participation.

### 2.2 Experimental procedure

#### 2.2.1 Task overview

We conducted two two-day experiments to investigate whether retrieval practice can attenuate stress-related memory inference impairments and examine the associated neural reactivation patterns. In both experiments, Day 1 consisted of the initial encoding phase, during which participants learned overlapping AB and BC image pairs. In Experiment 2 only, participants completed an additional retrieval-practice phase immediately after encoding, in which they freely recalled as many learned AB and BC pairs as possible without feedback. Experiment 1 did not include this retrieval-practice phase. Participants then left the laboratory and returned approximately 24 hours later for Day 2. No additional task or experimental manipulation was administered between Day 1 and Day 2, and participants were not given specific restrictions regarding activities such as alcohol consumption or sleep schedule during this interval. Detailed procedures are illustrated in Figure S1.

On Day 2, participants returned for baseline stress measurements, including heart rate recording and the Positive and Negative Affect Schedule (PANAS), and were then randomly assigned to the stress or control group. PANAS was administered again immediately after the stress or control procedure, with continuous heart rate monitoring. Approximately 20 minutes after stressor onset, participants began a three-alternative forced-choice A-C inference test while EEG was recorded. Memory for the directly learned AB and BC associations was subsequently tested for trial-by-trial behavioral analysis. Finally, participants completed a visual localizer task to train an EEG-based classifier to distinguish famous faces from famous buildings based on their neural response patterns. The classifier was then applied to the inference-phase EEG data to decode the reactivation of bridge-related category information. Behavioral performance and neural responses during A-C inference were the primary measures used to address our research questions.

#### 2.2.2 Stimuli

We employed two distinct sets of visual stimuli for this study. One set was used in the main memory task, comprising the encoding, inference, and retrieval phases, while the other was used for the localizer phase. The two sets were independent of each other, and all images are publicly available (see our statement on Data and code availability below) for future experiments or replication purposes.

Stimuli for the main memory task: The main memory task used 72 images divided into three categories: faces, buildings, and objects. Specifically, this set included 12 images of famous faces (e.g., Andy Lau), 12 images of iconic buildings (e.g., the Great Wall), and 48 images of everyday objects (e.g., an iPad). These images were selected to reflect items commonly encountered in the daily lives of Chinese college students. The stimuli were organized into 24 “ABC” memory triads, where A and C were always two distinct object images, and B was either a face or a building image. Stimuli for the localizer phase: The localizer phase included 108 images, evenly distributed across the same three categories (36 famous faces, 36 iconic buildings, and 36 everyday objects). The images of everyday objects were sourced from the THINGS database(Hebart et al., 2019), while images of famous faces and buildings were obtained from publicly available online sources. All images were resized to a resolution of 474 × 658 pixels and edited to remove distracting elements.

#### 2.2.3 Encoding phase

During the encoding phase, participants learned 48 memory pairs (AB and BC) derived from ABC triads, with each triad divided into two separate components (AB and BC) for independent encoding. Participants were explicitly instructed to learn the relationship between the two images displayed on the screen; however, no hints were provided regarding the potential to form a complete triad. Each encoding trial began with a fixation cross presented for 100 milliseconds, followed by the display of an AB or BC image pair for 2500 milliseconds. The next trial began immediately after the fixation cross was presented again. Each AB or BC pair was presented twice, with the positions of the images reversed between presentations. For example, if image A appeared on the left and image B on the right during the first presentation, their positions were swapped during the second presentation, with image B on the left and image A on the right. Notably, images A and C were never presented together during encoding, enabling a memory inference test on Day 2. To control the temporal interval between the presentation of AB and BC pairs, a mini-block design was employed, ensuring a trial interval of 1 to 4 trials between corresponding AB and BC pairs. The encoding phase consisted of two runs, with participants allowed to take breaks between runs to maintain focus throughout the task.

#### 2.2.4 Retrieval practice phase

In Experiment 2, a retrieval practice phase was conducted following the encoding phase. Participants were given a sheet of paper and were instructed to perform a free recall test of the previously studied pairs (both AB and BC). They were given two minutes to write down as many pairs as they could remember. No actual pictures were presented. Participants were instructed to write down the names of the pictures in each pair they recalled. At the end of the two-minute period, the experimenter collected the response sheets. Importantly, no corrective feedback was provided on their performance. This phase was not included in Experiment 1.

#### 2.2.5 Stress procedure

To ensure equivalent encoding contexts for both control and stress groups and to confine the stress effects to Day 2, all participants underwent the same encoding procedure on Day 1, without knowledge of their Day 2 group assignment (i.e., stress or control).

##### Stress induction

To induce acute stress, we employed a modified version of the Trier Social Stress Test (TSST), a well-established protocol for activating the hypothalamic-pituitary-adrenal (HPA) axis and the sympathetic nervous system(SNS)(Kirschbaum et al., 1993). The procedure was administered by a panel of three evaluators who maintained neutral expressions throughout. To amplify the socio-evaluative threat, participants were informed that their performance was being video-recorded for later analysis; no recordings were retained to ensure participant privacy. The stress protocol began with a 3-minute anticipation period, during which participants received instructions for an impromptu speech. This was followed by a 4-minute public speaking task, where participants were required to present themselves as candidates for a teaching certification, explaining how they would teach a collection of prose poems. Finally, participants engaged in an 8-minute mental arithmetic task. To increase the cognitive load and associated stress, we modified the standard TSST by replacing the serial subtraction task with challenging sequential two-digit multiplication problems (e.g., 56×47). Participants submitted answers via a keyboard within a 5-second window, and an audible alarm was triggered for incorrect or delayed responses. Pilot testing had confirmed that completing these calculations within the allotted time was exceedingly difficult.

##### Control condition

The control procedure was carefully designed to match the duration and environmental context of the TSST while eliminating its stressful components. This approach, which removes socio-evaluative threat and time pressure while retaining non-stressful cognitive tasks, is consistent with validated control procedures used in previous TSST studies (von Dawans et al., 2012; Buchanan et al., 2014; Smith et al., 2016). The procedure consisted of two parts. First, participants quietly read the same collection of prose poems for 7 minutes. Second, they completed a simple, self-paced arithmetic task (e.g., 2+3) for 8 minutes without time constraints or performance feedback. Participants were not disturbed, even for incorrect or delayed responses.

#### 2.2.6 Memory inference phase

The memory inference phase began approximately 20 minutes after the onset of the stressor. Before the formal test, we used slides and practice trials featuring completely unrelated images to explain the structure of the ABC task and to clarify that picture A and C were indirectly related through B. After confirming participants’ understanding of the inference task, they completed a three-alternative forced-choice memory inference phase. Each inference trial began with the presentation of a fixation point, displayed for a randomly selected duration (500 ms, 1000 ms, 1500 ms, or 2000 ms), followed by the memory cue (i.e., picture A) for 4000 ms. Participants were instructed to vividly retrieve and imagine the answer (i.e., picture C) during this period. This phase was the primary window of interest for EEG analyses, as successful inference required the recall of picture B (which we referred to as the “bridge information”) and picture C during this phase. Following the presentation of the memory cue, three candidate pictures were shown, and participants were required to select the correct answer within 3000 ms using the keyboard. One of the pictures was the correct answer (i.e., picture C), while the other two were distractor object images. These distractors had been presented during the Day 1 encoding phase but were not paired with the current memory cue A. This setup ensured that participants had to perform the full A-B, B-C inference to select the correct answer, rather than choosing based on familiarity alone (e.g., recognizing the picture from earlier encoding phase). If participants selected the correct option, the trial was labeled as correct. Additionally, we took further steps to account for potential guessing by analyzing trial-by-trial data from the retrieval phase. Specifically, we used trial-by-trial direct-memory performance to identify trials where the relevant AB and BC premise pairs were correctly answered, while acknowledging that this 2AFC measure cannot entirely rule out guessing.

#### 2.2.7 Direct memory retrieval phase

In addition to the A-C memory inference task, participants were also tested on their originally learned AB/BC pairs. We refer to this phase as direct memory retrieval to emphasize its distinction from the A-C inference, which involves both memory retrieval and inference. At the start of each direct memory retrieval trial, a fixation point was presented for a randomly selected duration (500 ms, 1000 ms, 1500 ms, or 2000 ms). Next, a memory cue (i.e., picture A or C) and two response options (one being the correct picture B) were presented simultaneously, with a 3000 ms window for participants to make a selection. Similar to the inference phase, the distractor image was a previously presented image from another AB or BC pair and belonged to the same category as the correct option. This task required direct recalling information from the encoding phase without engaging in memory inference. The direct retrieval phase consisted of 48 trials, testing all 24 AB and 24 BC pairs learned.

#### 2.2.8 Comparison between inference and direct retrieval phases

For the crucial A-C inference test, we used a three-alternative forced-choice (3-AFC) design where participants saw item A and had to choose the correct item C from three options. This design served two methodological purposes. First, it controlled for non-inferential response strategies. The two foil items were selected such that they were paired with other bridge elements (B items), with one foil’s bridge sharing the same stimulus category (face or building) as the correct triad’s bridge and the other foil’s bridge belonging to the different category. This forced participants to rely on the specific A-B-C association rather than a general categorical link. Second, the 3-AFC design reduces the probability of a correct guess to 33.3%, thereby increasing confidence that trials scored as ‘correct’ truly reflect successful inference. This is particularly important for the statistical power and validity of our subsequent EEG analyses contrasting successful and unsuccessful trials.

In contrast, the direct memory tests for A-B and B-C pairs employed a standard two-alternative forced-choice (2-AFC) design. Given that memory for these directly learned pairs was expected to be high, this format was used to assess memory for the directly learned pairs.

#### 2.2.9 Localizer phase

At the end of the experimental procedure, participants completed a visual category localizer task employing a 1-back paradigm with images from three distinct categories: famous buildings, famous faces (male and female), and common objects. EEG was used to measure brain activity. The stimuli for this task were unique and did not overlap with those used in the main memory tasks (encoding, inference, and retrieval phases). Each run of the task comprised nine mini-blocks, with three mini-blocks assigned to each image category, and each mini-block containing 12 trials. During each trial, an image was displayed for 1500 ms, followed by a 500ms inter-trial interval. Participants were instructed to respond quickly and accurately to indicate whether the current image was identical to the previous one. This task requirement served as a cover story to ensure participants remained engaged with the stimuli; however, their behavioral responses were not included in subsequent analyses. To prevent category-specific fatigue or habituation, the order of mini-blocks was pseudo-randomized, ensuring that two mini-blocks from the same category did not appear consecutively. The primary objective of the localizer task was to collect neural response patterns specific to each image category, which were later used to train a machine learning model for image classification based on brain activity (see *Decoding of EEG Signals* below).

#### 2.2.10 Physiological and Psychological measures of acute stress response

In Experiment 1, heart rate, as a physiological measure of acute stress, was continuously monitored using the BIOPAC MP160 system. In Experiment 2, heart rate was tracked with portable equipment, specifically the POLAR V2 system, which includes a Polar WearLink and a heart rate monitor. Heart rate was initially measured for 1 minute to establish a baseline and was continuously monitored throughout the subsequent tasks. Average heart rates were calculated for each experimental stage: a 3-minute preparation, a 4-minute public speaking, and an 8-minute arithmetic stage.

The Positive and Negative Affect Schedule (PANAS; Watson et al., 1988) was employed to assess participants’ current subjective affective states as a psychological measure of acute stress. This 20-item scale comprises 10 items for positive affects (e.g., “interested,” “excited”) and 10 items for negative affects (e.g., “nervous,” “scared”). Participants rated each item on a 5-point scale reflecting their current state, from 1 (very slightly or not at all) to 5 (extremely). Average scores for both positive and negative affect were then computed.

We acknowledge that cortisol, a key glucocorticoid hormone, is central to the physiological cascade initiated by acute stress and its subsequent effects on memory. However, this study was conducted during the COVID-19 pandemic under strict institutional safety protocols designed to minimize infection risk. As collecting saliva samples for cortisol assay would have posed an unnecessary risk of viral transmission to participants and research staff, we made the a priori decision to forgo this measure. The successful induction of stress was therefore confirmed using a combination of physiological (heart rate) and psychological (PANAS) metrics. Moreover, in a separate study conducted after COVID-19, we employed the same setup and stress-induction procedure and again observed significant increases in cortisol, further supporting the robustness of our protocol (Xu et al., 2025). We recommend that future research under normal conditions incorporates cortisol analysis to further elucidate the precise hormonal mechanisms.

### 2.3 EEG acquisition and preprocessing

EEG data were collected using Brain Products GmbH actiCHamp Plus amplifiers at a sampling rate of 1000 Hz. Continuous EEG was recorded from 64 electrodes, referenced to FCz, with an online band-pass filter of 0.1–100 Hz. Offline processing involved band-pass filtering between 0.1 and 30 Hz, down-sampling to 500 Hz, and re-referencing to the average of the left and right mastoids. To reduce line noise at 50 Hz, a band-stop filter was applied between 49 and 51 Hz. Subsequently, we visually inspected the ERP data and discarded any trials with residual ocular or other artifacts, such as those caused by movement. If an artifact was detected at any scalp site, data from all sites for that trial were excluded from further analysis. Interpolation was used to replace bad channels. Independent component analysis was performed to remove eye-blink and horizontal eye movement artifacts. After artifact rejection, an average of 23.88 ± 0.60 inference-phase EEG trials per participant were retained for subsequent decoding analyses. The continuous EEG signal was segmented into 2500 ms periods for the localizer task and 4500 ms for the inference task, each starting 500 ms before stimulus presentation. Segments were baseline-corrected against the mean voltage of the 500 ms pre-stimulus period. Behavioral data were used to label each trial with a true or false category, indicating whether the trial involved a famous building or a male or female famous face. The EEG data were preprocessed using the MNE-Python software package (Version 1.9.0), applying a band-pass filter from 0.1 to 30 Hz. Following filtering, the entire 0.1–30 Hz bandwidth was analyzed without further frequency decomposition.

### 2.4 Multivariate time-resolved decoding of EEG signals

#### 2.4.1 General overview

To investigate the neural reactivation of memory content, we performed a time-resolved multivariate decoding analysis on the EEG data using a Linear Discriminant Analysis (LDA) classifier, implemented with custom Python code adapted from the NeuroRA toolbox (Lu and Ku, 2020). For all decoding analyses, features were extracted from the preprocessed EEG signals using a sliding window approach. A 10 ms window (5 consecutive time points) was shifted in 10 ms increments across the epoch. The raw amplitude values from all 63 channels within each window were concatenated to form a feature vector (63 channels × 5 time points = 315 features) for each time step, preserving fine-grained temporal information.

To enhance the signal-to-noise ratio (SNR) of the EEG data before classifier training, we created trial aggregates (Isik et al., 2014; Grootswagers et al., 2017). For each participant and stimulus category (faces, buildings), we randomly sampled and averaged sets of 6 trials without replacement. This process was repeated until all trials for a given category were used, creating a balanced set of averaged trials (⌊N_trials / 6⌋ per category). To ensure the stability of our results, the entire decoding procedure, including the random sampling for trial averaging, was repeated 10 times, and the resulting accuracy maps were averaged across these iterations. We performed three distinct decoding analyses: (1) within-task classification to validate classifier performance, (2) cross-task classification to measure bridge element reactivation during inference, and (3) single-trial classification to link reactivation to behavioral success.

#### 2.4.2 Within-task classification

To validate the classifier’s ability to distinguish between face- and building-related neural patterns, we first conducted a cross-temporal decoding analysis exclusively on the localizer task data. Using the averaged trials, we employed a stratified 5-fold cross-validation procedure, repeated twice for stability. In each fold, the classifier was trained on 80% of the data and tested on the held-out 20%. The model trained at each time point was tested on all other time points, yielding a temporal generalization matrix for each participant. These matrices were then smoothed using a 2D moving average filter (5 × 5 time points, i.e., 20 ms × 20 ms) to highlight consistent decoding patterns.

#### 2.4.3 Cross-task classification

To test for the rapid neural reactivation of the category representation of bridge element B, we performed a cross-task classification. The LDA classifier was trained on the entirety of the averaged trial data from the localizer task (faces vs. buildings). This fixed decoder was then applied to EEG data from the memory inference task to classify neural patterns at each time point, despite no face or building images being present. This analysis yielded a temporal generalization matrix for each participant, where the training time corresponds to the localizer task and the testing time corresponds to the inference task. As a secondary analysis, the same decoder was applied to EEG data from the A-C choice phase to confirm the persistence of reactivation.

#### 2.4.4 Single-trial analysis of inference success

To directly link neural reactivation to memory inference performance, we applied the localizer-trained decoder to single trials from the inference phase, without trial averaging. We sorted trials based on whether the participant’s subsequent response was correct (successful inference) or incorrect (unsuccessful inference). This procedure generated separate temporal generalization matrices for successful and unsuccessful trials for each participant, allowing us to directly compare the strength of bridge element B reactivation as a function of behavioral outcome. On average, 13.61 ± 4.63 successful inference trials and 10.39 ± 4.63 unsuccessful inference trials per participant contributed to the single-trial decoding analysis.

### 2.5 Statistical analysis

#### 2.5.1 Validation of Acute Stress Induction

To assess the induction of acute stress, two-way mixed ANOVAs were performed on heart rate and subjective affect (separately for positive and negative affect), with group (stress vs. control) as a between-subjects factor and acquisition period as a within-subjects factor. We also defined three indices to represent individual differences in acute stress responses. For heart rate and negative affect ratings, we subtracted the baseline value from the post-induction value, with higher values indicating stronger stress responses. In contrast, for positive affect ratings, we subtracted the post-induction value from the baseline value, with a larger reduction indicating a stronger stress response.

#### 2.5.2 Analysis of Memory Inference Performance

To examine whether retrieval practice mitigated stress-induced impairments in memory inference, two-way ANOVAs were conducted on memory inference performance, with group (stress vs. control) and strategy (Experiment 1: no retrieval practice vs. retrieval practice) as between-subjects factors. Additionally, an independent t-test was performed in Experiment 1 to isolate the effect of acute stress on memory inference performance (i.e., whether acute stress impaired memory inference). Another independent t-test was conducted in Experiment 2 to compare the stress and control groups, as part of assessing whether retrieval practice was associated with attenuated stress-related inference impairment. Because a non-significant frequentist t-test does not establish evidence for the null hypothesis, we additionally conducted a Bayesian independent-samples t-test.

#### 2.5.3 Analyses of Decoding Accuracy Maps

We employed two approaches to analyze the generated decoding accuracy maps. The first analysis focused on determining whether mean classification accuracy exceeded the chance level of 50%. We applied permutation-based cluster-level correction to identify time windows with significant decoding accuracy, thereby suggesting the optimal training features and the timing of memory trace reactivation. Specifically, a cluster-based permutation test was applied to the 2D spatial data, which is well-suited for analyzing spatially correlated data, such as neuroimaging and EEG results. Initially, a one-sample t-test was performed for each spatial point in the 2D grid, comparing the results to chance level. A one-sided test with a p-value threshold of 0.05 identified significant points, which were assigned a value of 1, while non-significant points were marked as 0. Significant points were then grouped into clusters of adjacent significant points based on spatial connectivity. To assess cluster significance, a permutation procedure was applied, where the data were shuffled, and t-statistics were recalculated for 5000 iterations. The maximum t-statistic from each iteration was recorded, and clusters with observed t-statistics exceeding the 95th percentile of the permutation distribution were considered significant. The final output was a 2D matrix of binary values indicating the significance of each spatial point, ensuring robust control over Type I errors and identifying spatially contiguous regions of significant clusters.

#### 2.5.4 Optimal time window identification and validation

To compare neural reactivation strength between experimental conditions (e.g., Stress vs. Non-Stress) without incurring the problem of multiple comparisons across all time points, we focused our statistical analyses on an “optimal time window” that captured the peak of the reactivation signal. This window was derived from the pooled localizer data and validated using the following three-step procedure: (1) Identification of Subject-Level Peak Latency: We first identified the most robust time window for decoding in the classifier training data (the localizer task) for each participant individually. We performed a cluster-based permutation test on each participant’s time-series of decoding accuracy. We identified temporal clusters of significant decoding (defined as contiguous time points exceeding 50% accuracy for at least 10 ms). The “peak latency” for each participant was then defined as the center-of-mass of the cluster with the highest summed accuracy score. (2) Definition of the Group-Level Optimal Window: We then aggregated these peak latencies from all 136 participants. The group-level peak was the mean of this distribution, which was 197.46 ms (SD = 245.22 ms). Based on this, we defined the optimal training time window as 198±50 ms for our primary decoding analyses. (3) Cross-Task Validation: To ensure this time window was also representative of reactivation during the inference task, we performed a time-generalization MVPA. This analysis produced a 2D matrix showing classifier performance for every combination of training time (from the localizer task) and testing time (from the inference task). We applied the same center-of-mass procedure described in Step 1 to this cross-task decoding matrix to independently identify the peak reactivation time during inference. This validation yielded a peak latency of 246.01 ms (SD = 220.08 ms). This latency did not differ significantly from the one identified during model training (t(133)=1.66, *p*=0.10), indicating no significant temporal shift in the reactivation signature between tasks. This result validates our use of the fixed 198±50 ms window for all subsequent statistical comparisons of reactivation strength.

#### 2.5.5 Linking Neural Reactivation to Inference Success, Stress, and Retrieval Practice

Each participant generated three decoding accuracy matrices: (1) all trials, (2) trials with correct inferences (successful inferences), and (3) trials with incorrect inferences (unsuccessful inferences). Since each matrix is a high-dimensional structure containing over 200 × 450 accuracy outputs, we applied dimensionality reduction by averaging the values within the identified optimal training time window prior to further analysis. To quantify individual differences in memory reactivation within this window, we defined two indices representing the temporal and strength aspects of memory reactivation: *decodability* and *mean decoding accuracy*. Decodability was calculated as the percentage of time points with decoding accuracy above the 50% chance level within the specified time window. This index ranges from 0 to 100%, with higher values indicating that bridge-related category information was decodable at a greater proportion of time points during inference. In contrast, mean decoding accuracy was calculated by averaging decoding accuracy within the specified window, subtracting the 50% chance level, and multiplying by 100. This index ranges from −50 to 50 and represents decoding accuracy relative to chance in percentage points. Higher values indicate more accurate decoding of bridge-related category information.

To compare the neural indices of memory reactivation across experimental conditions, we defined the subsequent inference effect as the comparison of mean decoding accuracy and decodability between successful and unsuccessful inference trials, analogous to the subsequent memory effect, which compares neural activity between correctly and incorrectly recalled trials. For each participant, mean decoding accuracy and decodability were calculated separately for successful and unsuccessful inference trials. Paired-samples t-tests were then used to compare these within-participant measures across participants, with a two-tailed alpha level of p = 0.05. Additionally, we compared the two neural indices between Experiment 1 and Experiment 2 to examine the general effects of retrieval practice on memory reactivation, between stress and control participants in Experiment 1 to assess the effect of acute stress on memory reactivation, and between stress and control participants in Experiment 2 to investigate the protective effect of retrieval practice on memory reactivation under stress. For each decoding metric during inference, the within-experiment stress–control comparisons used two-sided independent-samples t-tests, with Holm correction applied across the two experiments separately for each metric. The directional hypotheses for the comparisons between experiments were informed by the behavioral findings; these comparisons were evaluated using one-tailed independent-samples t-tests and are interpreted as exploratory. The alpha level for these analyses was set at 0.05. Each specific hypothesis is outlined alongside its corresponding statistical test in the Results section.

### 2.6 Code availability

The custom code used in this study is available in https://osf.io/v54g8/?view_only=dd87e13d50b04c30b9ed839c58b80d95

### 2.7 Data availability

The data that support the findings of this study are in https://osf.io/v54g8/?view_only=dd87e13d50b04c30b9ed839c58b80d95

## Results

We conducted two independent EEG experiments with healthy young adults (total N = 136; Experiment 1: N = 68; Experiment 2: N = 68) to investigate how acute stress impacts memory inference and whether retrieval practice can mitigate such effects. In both experiments, participants encoded overlapping associative pairs (AB and BC) on Day 1, forming integrative ABC triads. On Day 2, they were tested on their ability to infer the indirect A-C relationship **(Figure 2)**. Experiment 1 investigated how acute stress, induced at the start of Day 2, affects inference performance and the underlying neural reactivation of the bridge element (B) compared to a non-stress control group. Experiment 2 assessed whether retrieval practice on the AB and BC pairs, implemented after encoding on Day 1, could buffer against these stress-induced impairments by modulating memory reactivation. The results are presented in five sections. First, we report manipulation checks confirming the efficacy of the stress induction at both psychological and physiological levels. Second, we present the effects of acute stress on memory inference accuracy and speed, and the mitigating role of retrieval practice. Third, we detail the results from our multivariate EEG decoding analyses of neural reactivation during inference. Fourth, we provide group comparisons of the time-resolved decoding outcomes to specify the effects of stress and retrieval practice. Finally, we examine whether the decoding results extend to the subsequent choice phase.

**Figure 2.**
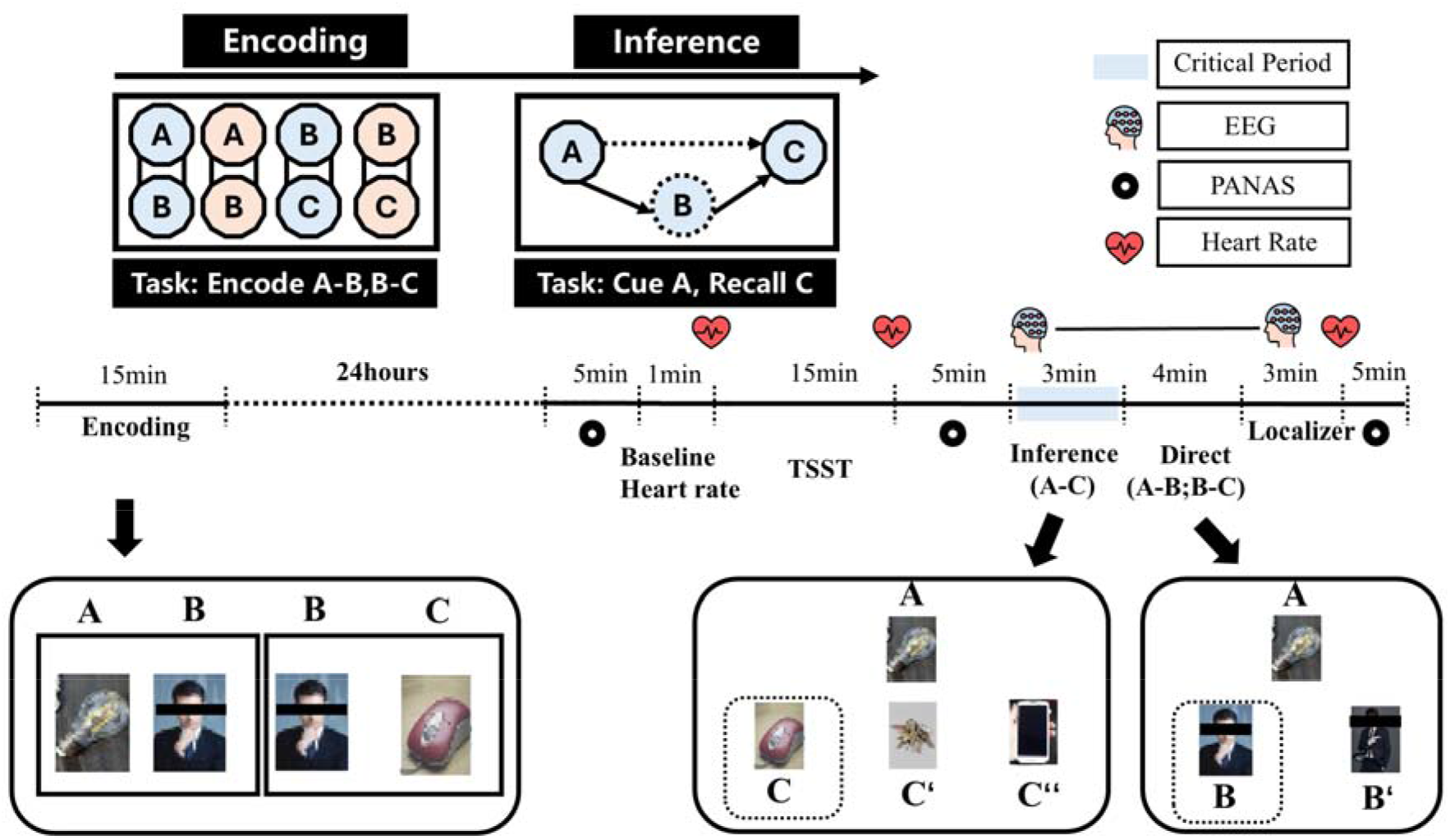
Task Overview. During encoding, participants learned overlapping AB and BC image pairs, forming interconnected triads (ABC). During inference, picture A was presented as a cue to recall picture C. Participants were not informed about the inference requirement during AB and BC encoding. The task comprised two sessions separated by approximately 24 hours. On Day 1, encoding lasted approximately 15 minutes, with two pictures presented side by side on the screen. On Day 2, participants underwent either a stress procedure (Trier Social Stress Test [TSST]) or a control procedure and then completed the memory tasks. Acute stress responses were assessed using heart rate measurements and the Positive and Negative Affect Schedule (PANAS). EEG data were recorded during the inference and localizer phases, with the inference phase serving as the primary period of interest. During the choice phase following inference, cue A was presented alongside the correct answer C and two lure images from other learned ABC triads. The original AB and BC associations were also tested to support trial-by-trial analyses of memory inference. The two experiments used the same procedure, except that Experiment 2 included retrieval practice on Day 1 immediately after encoding (see **Figure S1**).

### 3.1 Manipulation checks of stress induction

In both experiment1 and 2, we evaluated the effectiveness of acute stress induction using both physiological (heart rate in beats per minute, BPM) measures and psychological (self-reported Positive and Negative Affect Schedule, PANAS).

For physiological responses, we monitored heart rate throughout the stress induction procedure both experiments and during the memory inference phase in Experiment 1 and Experiment 2. Heart rate responses were quantified as the difference between the average during the Trier Social Stress Test (TSST) phase and the baseline phase (i.e., before stress induction). Independent t-tests showed significant higher heart rate elevation in the stress condition compared to the control condition in both experiments (exp1: t(66) = 9.38, *p* < .001, Cohen’s d = 2.27, 95% CI = [1.66, 2.88], **Figure 3A**; exp2: t(66) = 6.34, *p* < .001, Cohen’s d = 1.54, 95% CI = [1.00, 2.08], **FigureS2A**).

**Figure 3.**
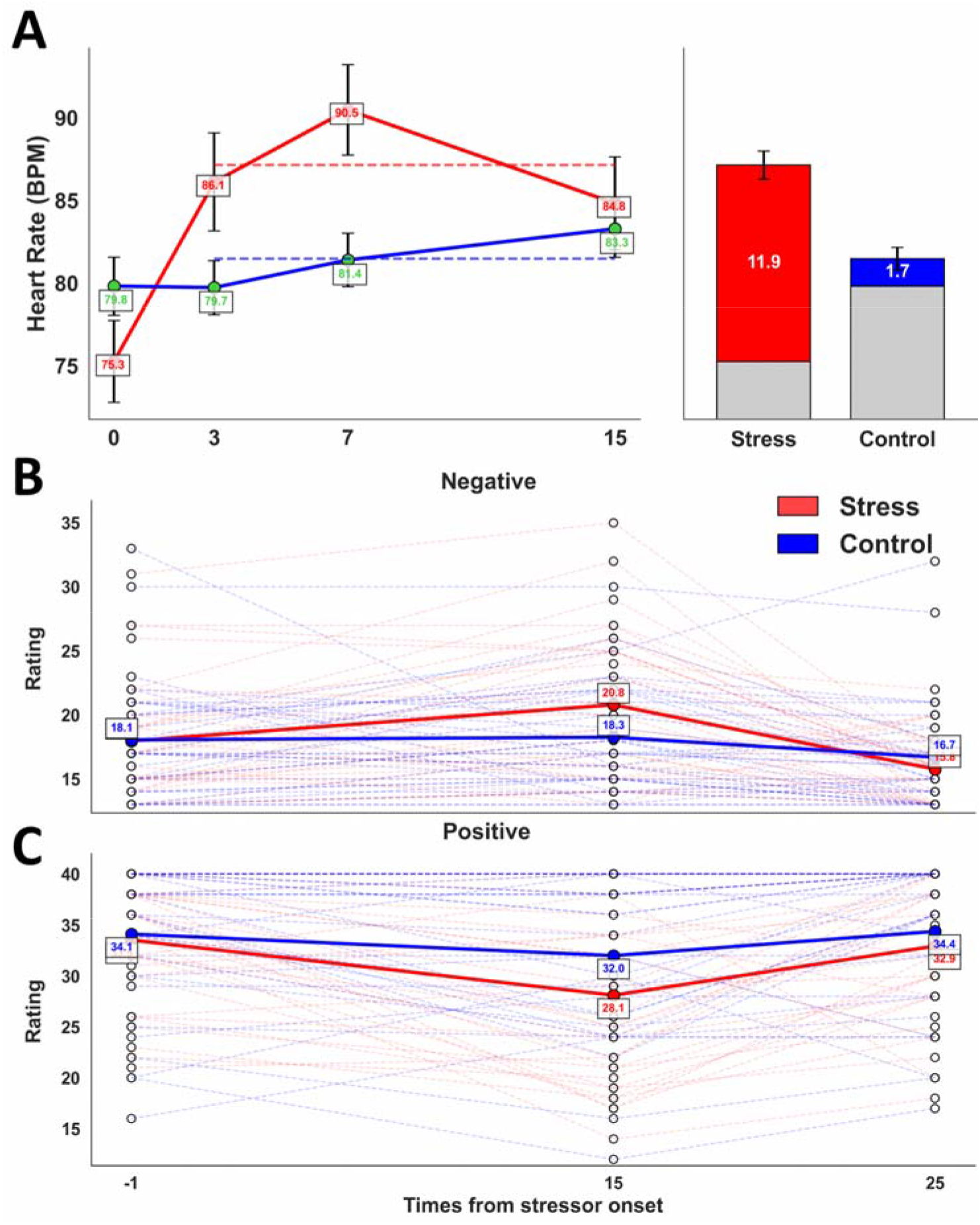
Manipulation Checks. **(A)** Heart-rate responses during the stress or control procedure in Experiment 1. The line plot shows mean heart rate in beats per minute (BPM) at baseline and during the three task periods of the stress/control procedure; error bars indicate SEM. The stacked bars summarize the heart-rate response relative to baseline. Grey segments indicate mean baseline heart rate, whereas colored segments indicate the mean increase from baseline to the average of the three task periods (red = stress group; blue = control group). Values shown inside the colored segments indicate the mean increase from baseline in BPM. Error bars on the stacked bars indicate SEM of this increase. The heart-rate response was significantly larger in the stress group (M = 11.9 BPM, SD = 4.95) than in the control group (M = 1.7 BPM, SD = 3.98). Group comparisons for the manipulation check were conducted on heart-rate change from baseline rather than raw BPM at each individual time point. **(B)** Stress significantly increased subjective ratings of negative valence. **(C)** Stress significantly decreased subjective ratings of positive valence. Note: Results from Experiment 1 are presented here. Similar stress responses were observed in Experiment 2, which are shown in **Figure S2**.

For psychological stress responses, we analyzed changes in PANAS scores. Differences between the second measurement (after stress induction) and the first measurement (before stress induction) served as indicators of stress responses for both negative and positive affect. Independent samples t-tests were conducted with group (stress vs. control) as the between-subject factor. In both experiments, the stress induction procedure led to significantly increased negative affect (exp1: t(66) = 2.33, *p* = 0.02, Cohen’s d = 0.57, 95% CI = [0.08, 1.05], **Figure 3B**; exp2: t(66) = 3.71, *p* < .001, Cohen’s d = 0.90, 95% CI = [0.40, 1.40], **Figure S2B**), and decreased positive affect in the stress group compared to the control group (exp1: t(66) = −2.15, *p* = 0.04, Cohen’s d = −0.52, 95% CI = [−1.01, −0.04], **Figure 3C**; exp2: t(66) = −2.46, *p* = 0.02, Cohen’s d = −0.60, 95% CI = [−1.08, −0.11], **Figure S2C**).

### 3.2 Memory performance

#### 3.2.1 Inference accuracy

In Experiment 1, we examined whether acute stress impaired memory inference (i.e., the ability to recall item C based on cue A in previously learned ABC triads). Memory inference performance was quantified using a normalized A-C inference index, calculated as A-C inference performance relative to direct AB/BC retrieval performance. This measure was used to account for individual differences in the availability of the directly learned premise pairs and does not represent raw 3AFC A-C accuracy.

Results revealed that acute stress significantly impaired memory inference compared to the control condition (t(66) = −2.33, *p* = 0.02, Cohen’s d = −0.56, 95% CI = [−1.05, −0.08], **Figure 4A left panel**).

**Figure 4.**
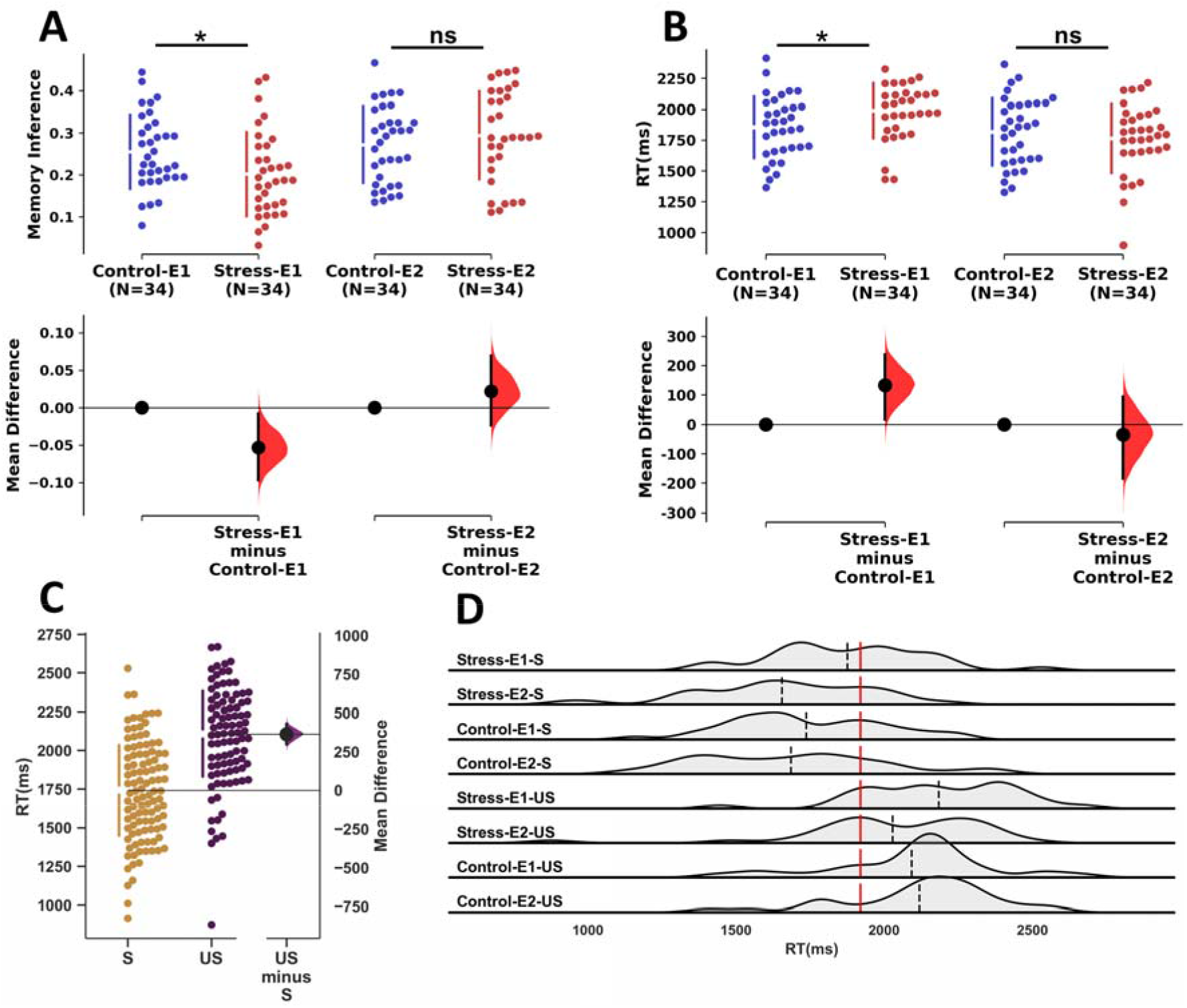
Memory inference performance. **(A)** Memory inference accuracy: Normalized A-C inference index across groups in Experiment 1 (Control-E1, Stress-E1) and Experiment 2 (Control-E2, Stress-E2). This index reflects A-C inference performance relative to direct AB/BC retrieval performance and therefore should not be interpreted as raw 3AFC accuracy or compared with the raw 3AFC chance level of 0.33. Acute stress significantly impaired inference accuracy in Experiment 1, whereas this impairment was attenuated in Experiment 2. **(B)** Response times (RTs) during the memory inference test. In Experiment 1, participants under acute stress exhibited significantly longer RTs compared to controls. No significant stress–control difference in RTs was detected in Experiment 2. **(C)** Comparison of RTs between successful (S) and unsuccessful (US) inference trials. Successful inferences were associated with significantly faster RTs compared to unsuccessful ones. **(D)** Density plots illustrating the distribution of RTs for successful (-S) and unsuccessful (-US) trials across different experimental conditions. Vertical lines indicate the mean of RTs.*p < 0.05.

Next, Experiment 2 tested whether retrieval practice could mitigate the stress-induced inference impairment reported in the Experiment1. Compared to Experiment 1, Experiment 2 included a retrieval practice phase immediately following the initial encoding of AB and BC pairs. On Day 2 participants underwent the same stress procedure and memory inference task as in Experiment 1. Unlike Experiment 1, no significant difference in memory inference performance was observed between the stress and control group (t(66) = 0.93, *p* = 0.35, Cohen’s d = 0.23, 95% CI = [−0.25, 0.70], **Figure 4A right panel**). A Bayesian independent-samples t-test provided weak evidence in favor of the null hypothesis, BF10 = 0.361 (BF01 = 2.77). Notably, the retrieval practice phase included only AB and BC pairs, with no AC inferences. Participants were informed that they would be tested on the practiced AB and BC pairs, and A-C associations were never directly practiced on Day 1. However, because we did not explicitly assess participants’ awareness of possible A-C links, we cannot fully rule out spontaneous awareness or integration of A-C associations during learning.

As a complementary cross-experiment analysis, we conducted a 2 (Stress: stress vs. control) × 2 (Retrieval Practice: no retrieval practice vs. retrieval practice) ANOVA on the primary A-C inference measure. This analysis revealed a significant main effect of retrieval practice (F(1, 132) = 11.10, *p* < .001, η^2^ = 0.07), no significant main effect of stress (F(1, 132) = 0.89, *p* = 0.35, η² = 0.01), and a significant stress-by-retrieval-practice interaction (F (1,132) = 5.24, *p* = 0.02, η^2^ = 0.04). Specifically, this interaction indicated that retrieval practice was associated with the attenuation of the stress-related inference impairment across experiments. We interpret this cross-experiment interaction cautiously because Experiment 2 was conducted as a follow-up experiment rather than as part of a single fully crossed factorial design.

An alternative explanation is that retrieval practice in Experiment 2 reduced stress levels during the memory inference phase, resulting in undamaged performance under stress. However, this is unlikely for two reasons: (1) between-group design ensured participants were unaware of retrieval practice manipulation in the other group, preventing confidence biases; and (2) retrieval practice and memory inference are distinct tasks, such that inference task-related stress would not be alleviated by prior retrieval practice on AB and BC. Additionally, we conducted data analyses to examine this alternative explanation. We analyzed the mean PANAS scores before and after the memory inference phase. We then compared these mean scores between the stress groups of Experiments 1 and 2 using two-sided independent-samples t-tests. Neither positive affect (t(66) = 1.16, p = .25, Cohen’s d = 0.28) nor negative affect (t(66) = 1.19, p = .24, Cohen’s d = 0.29) differed significantly between the two groups.

#### 3.2.2 Retrieval practice is associated with attenuated stress-related impairment in direct memory retrieval

We next examined whether stress had a broader impact on memory retrieval (i.e., for the individual premise pairs). Analysis of direct memory retrieval performance for A-B or B-C pairs showed that in Experiment 1, acute stress significantly impaired retrieval performance compared to the control group (t(66) = −2.16, *p* = 0.04, Cohen’s d = −0.52, 95% CI [−1.01, −0.04]). This indicates that acute stress did indeed produce a general deficit in retrieving the directed encoded memories. This deficit was not present in Experiment 2, in which participants had completed retrieval practice. We found no significant difference in direct memory performance between the stress and control groups (t(66) = −0.30, *p* = 0.77, Cohen’s d = −0.07, 95% CI [−0.55, 0.40]).

Specifically, a stress-induced reduction in A–C inference performance could partly reflect a broader impairment in retrieving the underlying premise associations (A–B and B–C). If the stress group recalled fewer premise pairs, fewer trials would enter the conditional A–C inference analysis, potentially confounding the observed group differences. To directly address this, we calculated the number of “foundation pairs” for each participant, defined as triads for which both the A-B and B-C premise pairs were correctly answered (Table S1). A comparison of these counts revealed no significant difference between the stress and control groups in either experiment. In Experiment 1, the groups had a similar number of foundation pairs available for inference (t(66) = −1.65, *p* = 0.10, Cohen’s d = −0.40, 95% CI [−0.88, 0.08]). Likewise, in Experiment 2, there was no significant difference between the groups (t(66) = −0.28, *p* = 0.78, Cohen’s d = −0.07, 95% CI [−0.54, 0.41]).

We further tested whether the cross-experiment stress-by-retrieval-practice pattern remained after controlling for the number of both-correct AB/BC triads. The stress-by-retrieval-practice interaction remained significant for the primary normalized inference index after including both-correct triad count as a covariate, b = 0.051, SE = 0.024, t = 2.17, p = .031, 95% CI [0.005, 0.098]. This analysis suggested that unequal numbers of foundation pairs may not fully explain the observed inference pattern.

#### 3.2.3 Retrieval practice enhanced direct memory retrieval but not baseline inference

To further probe the mechanism by which retrieval practice was associated with better inference under stress, we performed two additional analyses comparing performance across Experiment 1 (no retrieval practice) and Experiment 2 (with retrieval practice). First, we examined whether retrieval practice enhanced the memory for the directly encoded pairs. A t-test comparing the combined A-B or B-C accuracy across all participants from both experiments confirmed a significant benefit of retrieval practice, with higher accuracy in Experiment 2 than in Experiment 1 (t(134) = 2.41, *p* = 0.02, Cohen’s d = 0.41, 95% CI [0.07, 0.75]). This finding indicates that retrieval practice was associated with better memory for the premise pairs.

Second, we investigated whether this strengthening of direct memories led to a general improvement in inference, even in the absence of stress. To this end, we compared A-C inference accuracy between the control group of Experiment 2 and the control group of Experiment 1. This analysis revealed no significant difference in performance between the two groups (t(66) = 0.79, *p* = 0.43, Cohen’s d = 0.19, 95% CI [−0.28, 0.67]).

We additionally compared the two stress groups across experiments to directly assess whether retrieval practice was associated with better inference performance under stress. Participants in the retrieval-practice stress group showed higher A-C inference performance than those in the no-retrieval-practice stress group (t(66) = 3.74, p < .001, Cohen’s d = 0.91).

To further examine whether strengthened premise memory statistically accounted for the relationship between retrieval practice and A-C inference, we conducted an exploratory mediation analysis. Retrieval practice was coded as a binary predictor (No Retrieval Practice = 0, Retrieval Practice = 1), premise-memory performance was indexed by the number of correctly answered AB/BC direct-memory trials, and A-C inference was indexed by the primary inference measure. This analysis showed that retrieval practice was associated with better premise-memory performance (a = 2.13), and premise-memory performance was positively associated with A-C inference after controlling for retrieval practice (b = 0.012). The bootstrap indirect effect was significant (ab = 0.026, 95% CI [0.005, 0.049]), indicating that premise-memory performance partially accounted for the association between retrieval practice and later A-C inference. The direct effect of retrieval practice on A-C inference remained significant after accounting for premise-memory performance (c′ = 0.028, 95% CI [0.002, 0.055]). However, the coarse 2AFC measure administered after inference may not fully capture underlying premise-memory strength, which could contribute to the remaining association.

As an item-level robustness check, we fitted a mixed-effects model to trial-level A-C inference accuracy with fixed effects of stress, retrieval practice, and their interaction, and with random intercepts for participants and ABC triads. The stress-by-retrieval-practice interaction showed a positive trend consistent with the primary subject-level analysis, b = 0.116, SE = 0.061, z = 1.90, p = .058. The interaction estimate was in the same direction as the participant-level result but remained uncertain in this trial-level model.

#### 3.2.4 Inference speed

To investigate the effects of acute stress and retrieval practice on the speed of memory inference, we analyzed response times (RTs) during the A-C inference test. In Experiment 1, which did not include retrieval practice, participants under acute stress exhibited significantly longer RTs compared to the control group (t(66) = 2.32, *p* = 0.02, Cohen’s d = 0.56, 95% CI [0.08, 1.05]), indicating that acute stress slows memory inference. In contrast, no significant stress-related slowing was observed in Experiment 2 following retrieval practice, with no significant difference in RTs between the stress and control groups (t(66) = −0.81, p = 0.42, Cohen’s d = −0.20, 95% CI [−0.67, 0.28]). A combined analysis across both experiments revealed a significant main effect of retrieval practice (F(1, 132) = 8.69, *p* = 0.004, η^2^= 0.06), suggesting that retrieval practice enhances inference speed generally. The main effect of acute stress was not significant (F(1, 132) = 0.80, *p* = 0.37, η^2^= 0.01), reflecting the mitigation of stress effects in Experiment 2. Crucially, an integrated analysis revealed a significant interaction between acute stress and retrieval practice (F(1, 132) = 4.49, *p* = 0.04, η^2^= 0.03; **Figure 4B**), consistent with the interpretation that retrieval practice was associated with attenuation of stress-induced slowing across experiments. Across all conditions in both experiments, a clear speed-accuracy relationship was observed: successful inferences were associated with significantly faster RTs than incorrect ones (t(132) = −13.90, *p* < 0.001, Cohen’s d = −1.20, 95% CI [−1.43, −0.98]; **Figure 4C**). To probe the mechanism of the buffering effect, we examined whether retrieval practice differentially modulated RTs for successful versus unsuccessful inferences (**Figure 4D**). This analysis revealed that retrieval practice significantly accelerated RTs for successful inferences (t(134) = −2.78, *p* = 0.01, Cohen’s d = −0.48, 95% CI [−0.82, −0.14]) but had no significant effect on unsuccessful trials (t(131) = −1.37, p = 0.17, Cohen’s d = −0.24, 95% CI [−0.58, 0.10]). Retrieval practice was associated with faster responses on successful inference trials.

### 3.3 EEG results: Rapid neural reactivation of Bridge-related category information is associated with subsequent memory inference

To capture the neural dynamics underlying memory inference, we employed multivariate pattern analysis (MVPA) on time resolved EEG data. Our rationale was that if bridge information (i.e., picture B) is mentally reactivated during inference, then a classifier trained to discriminate between face and building images—using neural activity collected during an independent localizer phase—would successfully classify the category of the reactivated bridge information during inference, even though only object images (i.e., cue A and answer C) were presented (**Figure 5A**). We interpret this category-level reactivation as reflecting retrieval of bridge-related information that may contribute to constructing the indirect AC relationship. We used this neural decoding strategy as a means to capture underlying mental processes that are difficult to assess via self report and examined how this neural measure of memory reactivation supports memory inference. Following the principle of subsequent memory effect, we defined the subsequent inference effect contrast (i.e., successful vs. unsuccessful inference trials) to investigate whether neural reactivation of bridge-related category information predicted subsequent correct behavioral responses (**Figure 5B**). We present the EEG results in the following order: (1) within task classification, in which we trained and tested the model to classify the category of pictures on localizer phase data; (2) cross task classification, where the classifier was trained on localizer data and tested on inference phase data; and (3) analyses of the subsequent inference effect integrating outcomes of neural reactivation analyses and trial-level behavioral responses.

**Figure 5.**
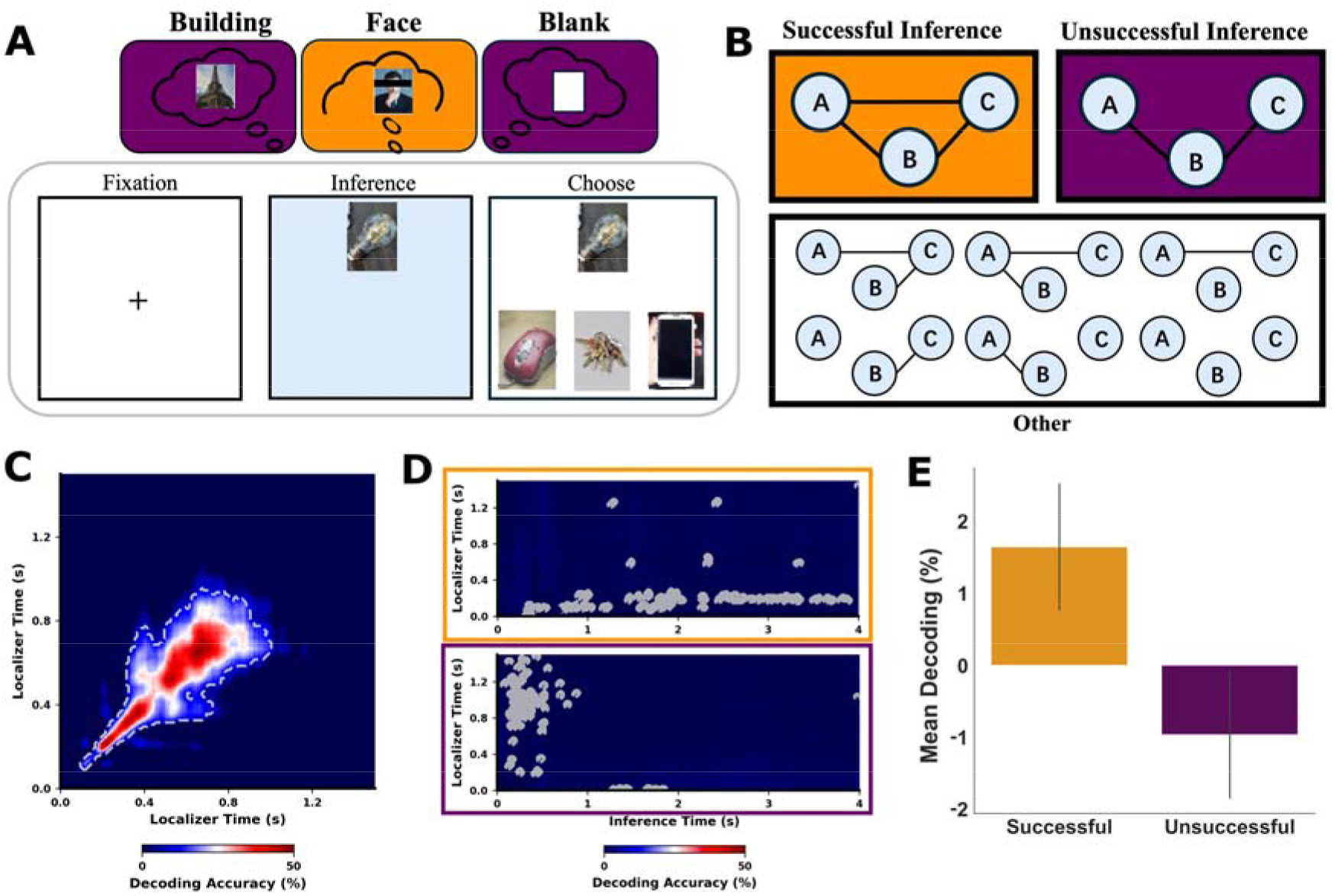
Bridge-related category information was reactivated to support subsequent successful inference. **(A)** Hypothesized reactivation of bridge information during the inference phase. To select the correct image-C based on cue-A, participants were hypothesized to mentally retrieve the bridge information-B (e.g., a face in the example trial, in orange). In some cases, the incorrect image (e.g., a building, in purple) might also be reactivated, or no image might be recalled during this phase. **(B)** Definition of successful and unsuccessful inference trials. In successful trials, both associations (A–B and B–C) were later answered correctly, allowing for the correct inference of the A–C relationship (Orange Box). In contrast, in unsuccessful trials, despite remembering A–B and B–C, participants failed to correctly infer the A–C relationship (Purple Box). Trials in which either A–B or B–C associations were not later answered correctly were excluded from the analysis (White Box). **(C)** Results of time-generalization analysis from within-task classification. EEG data from the localizer phase were used to train a machine learning model to classify faces and buildings based on localizer-phase EEG activity. **(D)** Results of time-generalization analysis from cross-task classification. EEG data from the localizer phase were used to train a machine learning model to classify faces and buildings using EEG data from the inference phase. Decoding accuracy matrices were computed separately for successful and unsuccessful inference trials (depicted in orange and purple, respectively). **(E)** Mean decoding accuracy within the optimal decoding time window, relative to chance, for successful and unsuccessful inference trials. Significantly above-chance decoding of bridge-related category information predicted subsequent successful inference.

We first validated the performance of our EEG multivariate classifier. We trained and tested the classifier on an independent localizer dataset using a cross-validation procedure (see **Methods**). The classifier was highly effective at distinguishing the stimulus categories (e.g., faces vs. buildings) from the EEG data (**Figure 5C**). The mean peak decoding accuracy across participants was 77.41% (SD=8.38%, 95% CI [75.99%,78.83%]), with individual peak accuracies ranging from 56.20% to 95.60%. On average, decoding accuracy was significantly above chance for a duration of 481.71 ms (SD=195.23 ms). The mean latency of the peak decoding accuracy was 458.22 ms (SD=192.40 ms) post-stimulus onset.

Next, to investigate whether and when the visual category (i.e., face or building) of bridge information (B) can be decoded from brain activity during an A–C inference trial, we applied cross task classification to the time resolved EEG data collected during the inference phase. The classifier was trained on neural activity elicited when participants viewed pictures from specific categories and then applied to the time window during which memory cue A was presented, prior to the selection of answer C. We hypothesized that the category information of bridge information (B) would be reactivated, despite the absence of face or building stimuli during this phase. To examine the temporal dynamics of rapid neural reactivation at the single trial level, we conducted a series of time resolved decoding analyses. Linear discriminant analysis (LDA; see Methods) was employed to classify visual category information based on the EEG topography at each time point following the presentation of memory cue A. Two primary cross task classification clusters were identified in the decoding accuracy maps. The first cluster extended from 120 to 290 ms during the localizer phase and from 740 to 3890 ms during the inference phase. The second cluster spanned 1310 to 1500 ms during the localizer phase and comprised two distinct intervals during the inference phase: 90 to 1500 ms and 2680 to 4000 ms (**Figure S3A**).

Lastly, to validate the behavioral relevance of the identified neural reactivation of bridge information (B), we examined whether this reactivation predicted subsequent successful inference (i.e., Subsequent Inference Effect). Each trial was labeled as either successful or unsuccessful, and we anticipated that rapid neural reactivation would be more pronounced in successful trials compared to unsuccessful ones. For each participant, two decoding matrices were generated: one for successful inference and one for unsuccessful inference. Two participants were excluded from the Subsequent Inference Effect analyses because two types of trials could not be defined for them, resulting in a final sample of 134 participants. When focusing on neural activity preceding successful inference, we observed significant cross task classification, ranging from 0 ms to 250 ms in the localizer phase and from 310 ms to 3890 ms in the inference phase (**Figure 5D, upper orange panel**). In contrast, when applying the classifier to neural activity preceding unsuccessful inference, we observed significant cross task classification, ranging from 200 ms to 1500 ms in the localizer phase and from 70 ms to 890 ms in the inference phase (**Figure 5D, lower purple panel)**. Moreover, we identified the optimal time window to capture memory reactivation (See Methods for details), and then averaged each participant’s 2D maps across time points within that time window and adjusting by subtracting the 50% chance level, thereby setting 0% as the new baseline. An initial paired t-test revealed that the classifier decoded reactivated bridge information (B) more accurately prior to successful inference than unsuccessful inference (t(133) = 2.11, *p* = 0.037, Cohen’s d = 0.182, 95% CI = [0.16, 5.05]; **Figure 5E**). Follow-up tests indicated that the category of bridge information was significantly reactivated above chance levels prior to a successful inference response (t(133) = 1.87, *p* = 0.031, one-tailed, Cohen’s d = 0.162). Conversely, for unsuccessful inference trials, decoding accuracy did not significantly differ from chance (t(133) = −1.09, *p*= 0.862, one-tailed, Cohen’s d = –0.094). These findings underscore the behavioral relevance of the detected reactivation of bridge information during the inference phase.

### 3.4 EEG results: Acute stress disrupts reactivated neural representations of bridge-related category information, but retrieval practice is associated with attenuation of this disruption

The memory inference phase began approximately 20 minutes after the onset of the stressor. Behavioral data indicate that acute stress impairs memory inferences in humans. To investigate how acute stress influences the neural mechanisms underlying the flexible use of encoded memories to infer A–C associations, we employed rapid neural reactivation of bridge information as a neural lens. We used two neural indices to capture the dynamics of memory reactivation: *Decodability* and *Mean Decoding Accuracy* (see Methods for details). *Decodability* reflects the temporal consistency of memory reactivation, with higher scores suggesting sustained activation during the inference phase. In contrast, *Mean Decoding Accuracy* quantifies the strength of memory reactivation by averaging the accuracy of decoding bridging information from brain activity, adjusted for chance levels; higher values indicate stronger retrieval of memory representations.

Initially, we examined whether stress reduced the reactivation of bridge-related category information during inference. In Experiment 1, we compared *Decodability* and *Mean Decoding Accuracy* between stress (Stress-E1) and control participants (Control-E1). Relative to the control condition, acute stress significantly reduced the reactivation of bridge information, as reflected in lower *Decodability* (t(66)=-2.19, *p_Holm* =.032; Cohen’s d=−0.53) and decreased Mean Decoding Accuracy (t(66)=−2.48, *p_Holm=*.032, Cohen’s d=−0.60). To further support the notion that acute stress disrupts neural reactivation of bridge information, we correlated individual differences in acute stress responses (i.e., changes in HR and PA) with *Decodability* and *Mean Decoding Accuracy* measures. To avoid interpretational issues with pooled data, we conducted correlation analyses separately for each experiment and found no significant correlations. In Experiment 1, we found no correlation for either decodability (Stress: r = −0.12, *p* = 0.50; Control: r = −0.19, *p* = 0.28) or mean decoding accuracy (Stress: r = −0.05, *p* = 0.76; Control: r = −0.23, *p* = 0.18). This pattern was replicated in Experiment 2 for both decodability (Stress: r = −0.07, *p* = 0.76; Control:r = −0.03, *p* = 0.89) and mean decoding accuracy (Stress: r = −0.04, *p* = 0.82; Control: r = 0.12, *p* = 0.48).

In Experiment 1, participants under stress were less likely to infer correct A–C associations. However, behavioral data suggest that retrieval practice of AB and BC associations was associated with attenuated stress-induced impairment and better overall inference performance. If neural reactivation of bridge information contributes to successful inference and reflects the impact of stress, then it should be modulated by retrieval practice.

As an exploratory analysis informed by the behavioral findings, we compared the decoded neural dynamics of bridge-related category information during inference between Experiment 2 and Experiment 1, collapsing across stress conditions. Participants in Experiment 2 showed significantly stronger reactivation of bridge information than those in Experiment 1, as reflected in higher Decodability (t(134) = 1.79, *p* = .037, one-sided, Cohen’s d = 0.30) and higher Mean Decoding Accuracy (t(134) = 1.74, *p* = .042, one-sided, Cohen’s d = 0.29).

Subsequently, we examined participants in Experiment 2 who all underwent retrieval practice but received either stress (Stress-E2) or control (Control-E2) manipulations prior to inference, to elucidate why the memory inference process is resilient to stress following retrieval practice. We observed that participants in Experiment 2 who received stress not only avoided the stress-induced impairment in reactivation but also exhibited significantly stronger reactivation of bridge information compared to those receiving control manipulations, as reflected in both higher Decodability (t(66) = 2.75, *p_Holm =* .015, Cohen’s d = 0.66) and higher Mean Decoding Accuracy (t(66) = 2.22, *p_Holm=*.032, Cohen’s d = 0.53).

Finally, as a complementary analysis, we combined data from both experiments to formally test the interaction between acute stress and retrieval practice, acknowledging that Experiment 2 was designed after the results of Experiment 1 were known. For decodability of bridge-related category information, this analysis revealed a significant acute stress × retrieval practice interaction (F(1, 132) = 12.32, p < .001, ηp² = .085; Figure 6A). Holm-corrected simple-effect tests showed lower decodability in the stress than in the control group in Experiment 1 (adjusted p = .032), whereas decodability was higher in the stress group in Experiment 2 (adjusted p = .015). A similar pattern emerged for Mean Decoding Accuracy: the 2 × 2 ANOVA revealed a significant acute stress × retrieval practice interaction (F(1, 132) = 10.77, p = .001, ηp² = .075; Figure 6B), consistent with an association between retrieval practice and altered stress effects on inference-related decoding.

**Figure 6.**
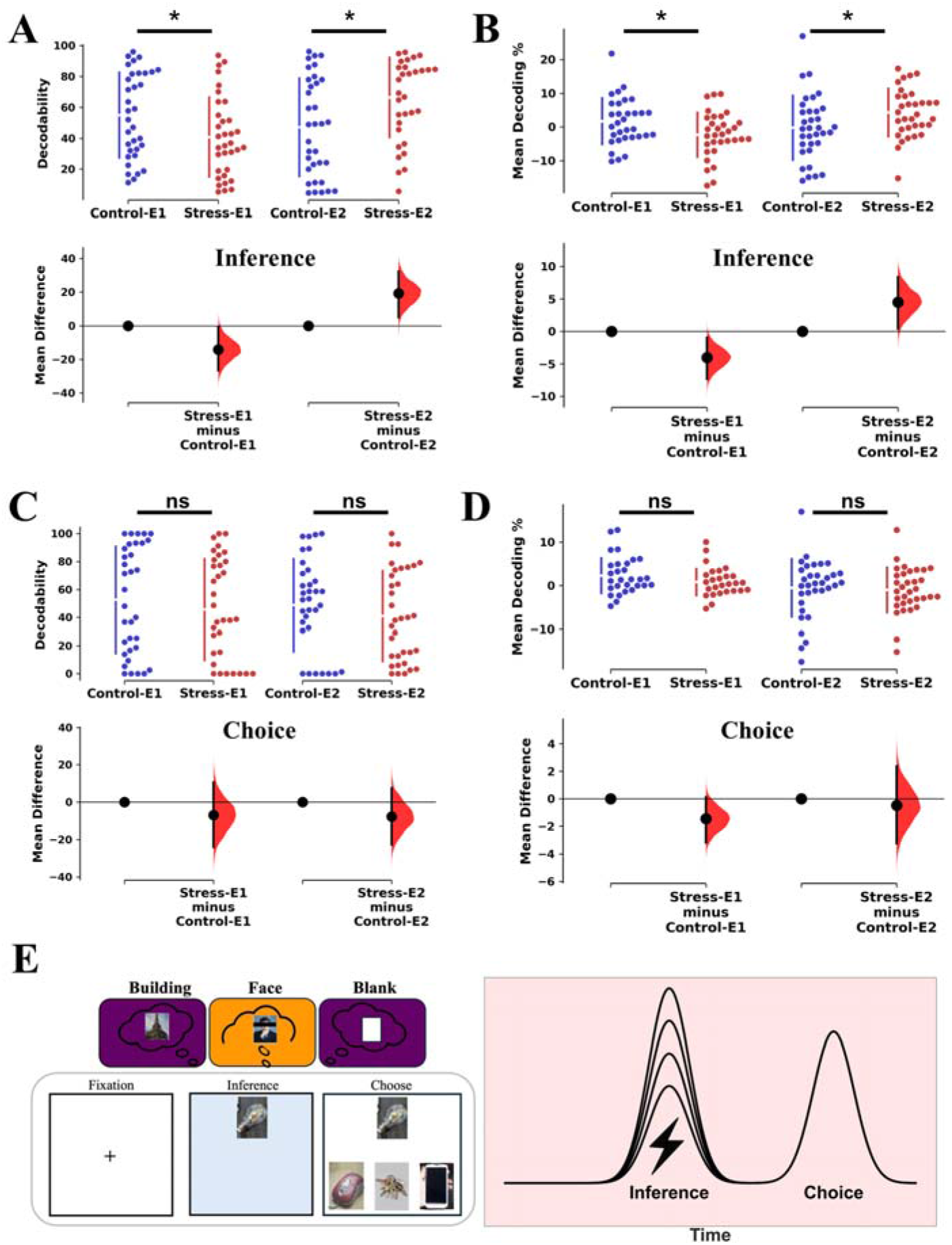
Neural reactivation under acute stress and after retrieval practice. Decodability **(A**) and Mean Decoding Accuracy (**B**) of the bridge element during the Inference Stage. Acute stress reduced neural reactivation in Experiment 1 (No-Retrieval Practice) but enhanced it in Experiment 2 (Retrieval Practice). \**P*corr < 0.05. Decodability (**C**) and Mean Decoding Accuracy (**D**) of bridge-related category information during the Choice Stage. In contrast to the inference stage, choice-phase reactivation showed no significant interaction between acute stress and retrieval practice. (E) Schematic of the trial structure, illustrating the distinct inference and choice phases analyzed. These findings indicate that the modulatory effects of stress and retrieval practice on bridge-related category element reactivation are confined to the inference process itself.

### 3.5 EEG results: Neural reactivation of bridge element during the choice phase predicts successful inference but is unaffected by stress or retrieval practice

As a complementary control analysis, we examined the choice phase to determine whether bridge-related category reactivation was specific to the initial inference period or also present during the later response-selection period. This analysis addressed whether the stress- and retrieval-practice effects observed during the cue/inference period generalized to the subsequent choice phase. Because participants selected their response during this 3-s window, this analysis was interpreted cautiously and was not used as the primary neural test of inference-related reactivation. Applying the same multivariate decoder to the choice phase—during which participants viewed the cue (A) alongside the potential target (C)—revealed significant reactivation of the bridge element across all conditions (**FigureS3B**). Cluster-based permutation tests on the decoding time course identified three significant temporal clusters: two early clusters from 330–480 ms and 580–1110 (*p*<.05) and a more sustained later cluster from 2030–2660 ms (*p*<.05). To quantify this effect, we extracted the mean decoding accuracy within these time windows and investigated its functional relevance: this choice-phase reactivation of the bridge element was significantly greater on successfully inferred trials compared to unsuccessful trials (t(133) = 2.92, *p* = 0.004, Cohen’s d = 0.25, 95% CI =[0.56, 2.94]).

Crucially, in a key divergence from the initial inference period, this choice-phase reactivation was not modulated by interaction between acute stress and retrieval practice. We formally tested this using two complementary metrics. First, a 2 (Stress: Stress vs. No-Stress) × 2 (Session: Retrieval Practice vs. No-Retrieval Practice) ANOVA on overall decodability yielded no significant interaction between Stress and Retrieval Practice (F(1,132)=0.01, p=0.94, η ^2^<0.001; **Figure 6C**). A parallel ANOVA on mean decoding accuracy yielded also no significant interaction (F(1,132) = 0.35, *p* = 0.56, η_p_^2^ = 0.003; **Figure 6D**). In sum, while bridge element reactivation occurs during the choice phase and predicts performance, this neural signal—unlike the rapid reactivation during the initial inference stage—showed no interaction between Stress and Retrieval Practice.

## Discussion

Our study shows that acute stress is associated with impaired memory-based inference and reduced rapid bridge-related reactivation. We demonstrate that retrieval practice was associated with attenuated stress-related impairment in both behavioral performance and underlying neural reactivation. Using multivariate decoding of EEG data, we identified the rapid reactivation of bridge-related category information as a critical predictor of successful inference (A-C). This reactivation process was attenuated by stress, whereas retrieval practice was associated with stronger reactivation under stress. These findings provide a novel neuroscientific account of how memory inference is accomplished, clarify its vulnerability to stress, and highlight a powerful behavioral strategy for building stress resilience.

A central contribution of our work is the identification of rapid bridge-related category reactivation associated with successful memory-based inference. While prior neuroimaging research has focused on the roles of the hippocampus and prefrontal cortex in integrating memories during encoding (Zeithamova and Preston, 2010; Zeithamova et al., 2012; Backus et al., 2016; Morton et al., 2020; Schlichting et al., 2021), our work elucidates the neural dynamics of the inference phase itself. We reveal that successful A-C inference is preceded by a rapid neural reactivation of the bridge-related category information. This extends prior decoding studies of deliberate retrieval or imagery (Polyn et al., 2005; Chadwick et al., 2010; Xue, 2018; Dijkstra et al., 2019), as the reactivation observed here may reflect goal-relevant retrieval processes contributing to the linking of discrete memories. One consideration is that participants were informed of the A-C link via B on Day 2, which could encourage a deliberate reactivation. However, as these instructions were identical across all groups, they cannot account for the observed interaction between stress and retrieval practice, nor for the trial-by-trial relationship between reactivation strength and inference success. Future research could employ more implicit paradigms to disentangle the contributions of spontaneous versus goal-directed reactivation to memory inference.

Our results extend the literature on stress-induced memory retrieval deficits (de Quervain et al., 1998; Kuhlmann et al., 2005; Smeets, 2011; Shields et al., 2017; Wolf, 2017) to the domain of retrieval-dependent, more complex and constructive cognition. Consistent with the theoretical framework proposing that stress promotes a shift away from flexible, hippocampus-dependent cognitive strategies toward more rigid, striatum-based systems (Knowlton et al., 1996; Kluen et al., 2017b, 2017a; Raio et al., 2017; Vogel et al., 2018; Schwabe et al., 2022; Zerbes et al., 2022), we observed that acute stress impaired the retrieval of AB and BC associations and, in parallel, reduced the accuracy and speed of AC inference. While EEG lacks the spatial resolution to definitively dissociate hippocampal from striatal contributions, our data provide temporal evidence that stress is associated with reduced rapid reactivation of bridge-related category information during inference. We interpret this category-level reactivation as potentially reflecting retrieval of shared mnemonic information linking related events, which may contribute to constructing the indirect AC relationship. An alternative interpretation is that stress impairs the retrieval of A-C associations that were spontaneously formed during encoding (Zhou et al., 2023). However, the trial-by-trial relationship between bridge-related category reactivation and subsequent inference success, together with the behavioral and neural differences associated with retrieval practice, may reflect a contribution of online retrieval to inference. Importantly, because stress also impaired retrieval of AB and BC pairs, our data cannot fully disentangle a general retrieval deficit from a more specific disruption of inferential integration. Our findings therefore suggest that stress-related inference impairment may involve reduced retrieval of relevant memory information, whereas retrieval practice was associated with stronger category-level reactivation and better inference performance under stress.

Our study links retrieval practice (RP) to attenuation of the detrimental effects of acute stress on memory inference. Building on extensive evidence that RP strengthens memory (Karpicke and Blunt, 2011; Roediger and Butler, 2011; Liu et al., 2020; Ye et al., 2020; Yang et al., 2021; Zhuang et al., 2021; Carpenter et al., 2022), we found that practicing the AB and BC pairs was associated with reduced stress-related impairment in novel A-C inference. This generalization of the testing effect to a novel, integrative inference task contrasts with prior findings where its benefits are often contingent on congruence between practice and test formats (Smith et al., 2016; Brown et al., 2020).

How does retrieval practice confer this stress resilience at the neural level? We propose that practicing retrieval of the AB and BC pairs serves to “tag” the entire ABC memory structure for subsequent offline consolidation (Dragoi and Tonegawa, 2011, 2013), potentially through co-reactivation during practice or subsequent sleep (Lutz et al., 2024). Our results suggest that retrieval practice was associated with stronger premise memories and attenuated stress-related inference impairment. These benefits may reflect strengthened AB and BC associations, potentially accompanied by more effective online integration, although A-C associations formed before the test may also contribute.

Intriguingly, retrieval practice was associated not only with attenuated stress-related reductions in bridge-related category information reactivation but also with a stronger reactivation signal under stress. The decoding strength for bridge-related category information was highest in the RP-Stress group, significantly surpassing that of the RP-No Stress group. We interpret this heightened reactivation as potentially reflecting a compensatory neural mechanism. Under acute stress, stronger reactivation of bridge-related category information may contribute to inference performance. This interpretation aligns with broader theories of cognitive effort, which posit that additional neural resources are recruited to preserve performance in situations such as aging (Reuter-Lorenz and Cappell, 2008), facing increased task difficulty (Gray et al., 2003), or physiological challenges such as stress (Hermans et al., 2014).

To supplement our primary multivariate decoding results from the inference phase, we conducted an additional analysis of the choice phase. We applied the same decoder to the choice phase following inference. While this analysis also detected bridge-related category reactivation that predicted subsequent choice accuracy, this reactivation showed no significant stress-by-experiment interaction. We propose that this later reactivation may be cognitively distinct from that observed during initial inference, potentially reflecting post-inferential processes such as choice confirmation. This pattern suggests that stress-related differences in category reactivation may depend on the task phase. This dissociation suggests the time-specific effect of acute stress: it selectively disrupts neural reactivation during the cognitively demanding period of memory inference.

Our study is not without limitations, which highlight key directions for future investigation. First, although we used well-established stimulus categories (i.e., faces, scenes, objects), the generalizability of our findings could be further tested using materials with greater ecological validity or semantic complexity. Second, the limited spatial resolution of EEG prevented us from identifying the precise anatomical sources of the reactivation signals. Future studies employing methods with higher spatial resolution are needed to elucidate the network-level interactions between the hippocampus, prefrontal cortex, and other relevant regions during stress-impaired inference. Third, while our findings associate retrieval practice with attenuated stress-related inference impairment, a valuable direction for future research would be to include a restudy control group. Such a design would allow for a precise dissociation of the cognitive mechanisms attributable to active retrieval versus those stemming from mere re-exposure to the material, further clarifying the unique role of retrieval in building stress-resilient inferential memories. Nonetheless, a substantial body of literature has established the superiority of retrieval practice over restudy for enhancing long-term retention (Roediger and Butler, 2011; Roediger and Abel, 2022) and building stress-resilient memories (Smith et al., 2016), providing a strong rationale for our interpretation. In addition, the post-inference direct-memory test used a two-alternative forced-choice format, which provides only a coarse measure of premise-memory strength and allows some correct responses to arise from guessing. Therefore, restricting analyses to trials in which both AB and BC pairs were later answered correctly does not guarantee that those trials were supported by strong associative memories. Future studies using cued recall, confidence ratings, or continuous measures of premise-memory strength would provide a more sensitive test of how premise memory supports later inference. A further limitation is that retrieval practice was implemented in a separate follow-up experiment rather than within a single fully crossed 2 × 2 factorial design. Although the two experiments were closely matched in recruitment procedures, participant population, laboratory setting, EEG system, task structure, and experimenter team, cohort-related confounds cannot be fully ruled out. Therefore, the cross-experiment stress-by-retrieval-practice analyses should be interpreted cautiously. Future studies using a fully crossed factorial design are needed to provide a more rigorous test of the interaction between retrieval practice and acute stress. Some directional neural comparisons were informed by the behavioral findings and evaluated using one-tailed tests. These comparisons should therefore be interpreted cautiously. Because retrieval practice was implemented as a brief free-recall task, we could not reliably link retrieval success for individual AB/BC associations to later A-C inference trials. Therefore, the present data do not allow a direct comparison of later inference performance for successfully versus unsuccessfully practiced items. The item-level mixed-effects analysis served as a robustness check, but its interaction estimate remained uncertain. Future studies using trial-by-trial cued retrieval practice will be needed to test item-specific retrieval-practice benefits more directly. Finally, pandemic-related safety protocols precluded cortisol collection, and the available measurements cannot distinguish the contributions of glucocorticoid and adrenergic activity to the observed effects. Future studies should integrate hormonal measures to provide a more comprehensive characterization of the stress response.

In conclusion, our study shows that acute stress impairs memory inference and is associated with reduced rapid reactivation of bridge-related category information. This reactivation was associated with subsequent inference success and may contribute to memory-based inference under stress. Critically, we show that retrieval practice was associated with attenuation of stress-induced inference impairment across two closely matched experiments. This protection is associated with the preservation and even enhancement of the underlying rapid reactivation signals. These findings highlight retrieval practice as a potential strategy associated with more resilient memory-based inference under stress.

## Acknowledgments

This study was supported by the National Natural Science Foundation of China (grant No. 32300879 and W2421004 to W.L.), Humanities and Social Sciences Fund, Ministry of Education (grant No. 22YJCZH109 to W.L.).

## Supplementary

**Figure S1.**
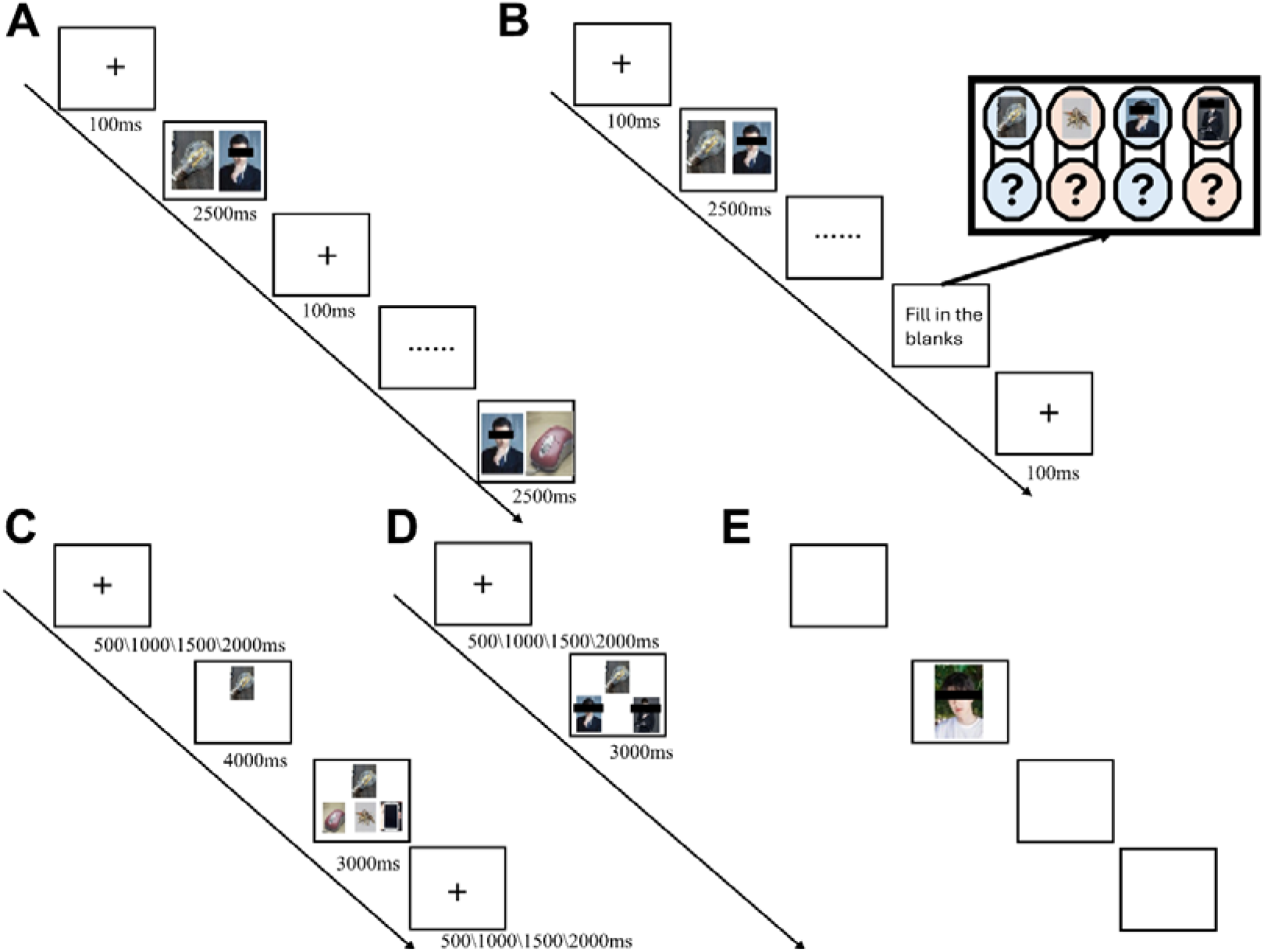
Experimental procedures. **(A) Encoding phase.** Each trial began with a fixation cross (100 ms), followed by the presentation of an AB or BC image pair for 2500 ms. Each pair was presented twice, with the spatial positions reversed in the second presentation. **(B) Retrieval practice phase.** Following the encoding phase, participants were instructed to freely recall and fill in the blanks for the learned memory pairs (AB and BC) using paper and pen within a two-minute time limit. **(C) Memory inference phase.** Each trial started with a fixation point presented for a jittered duration (500, 1000, 1500, or 2000 ms), followed by a memory cue (Item A) for 4000 ms. Subsequently, three options were displayed, and participants were required to select the correct target (Item C) within 3000 ms. Distractors were images from the encoding phase (Day 1) that were not paired with the current cue. **(D) Direct memory retrieval phase.** After a jittered fixation, a cue (Item A or C) and two options (one being the correct Item B) were presented simultaneously. Participants had 3000 ms to make a selection. The distractor was a previously learned image from the same category as the correct option. **(E) Localizer phase.** Images were presented for 1500 ms with a 500 ms inter-trial interval. Participants performed a 1-back task, indicating whether the current image was identical to the preceding one.

**Figure S2.**
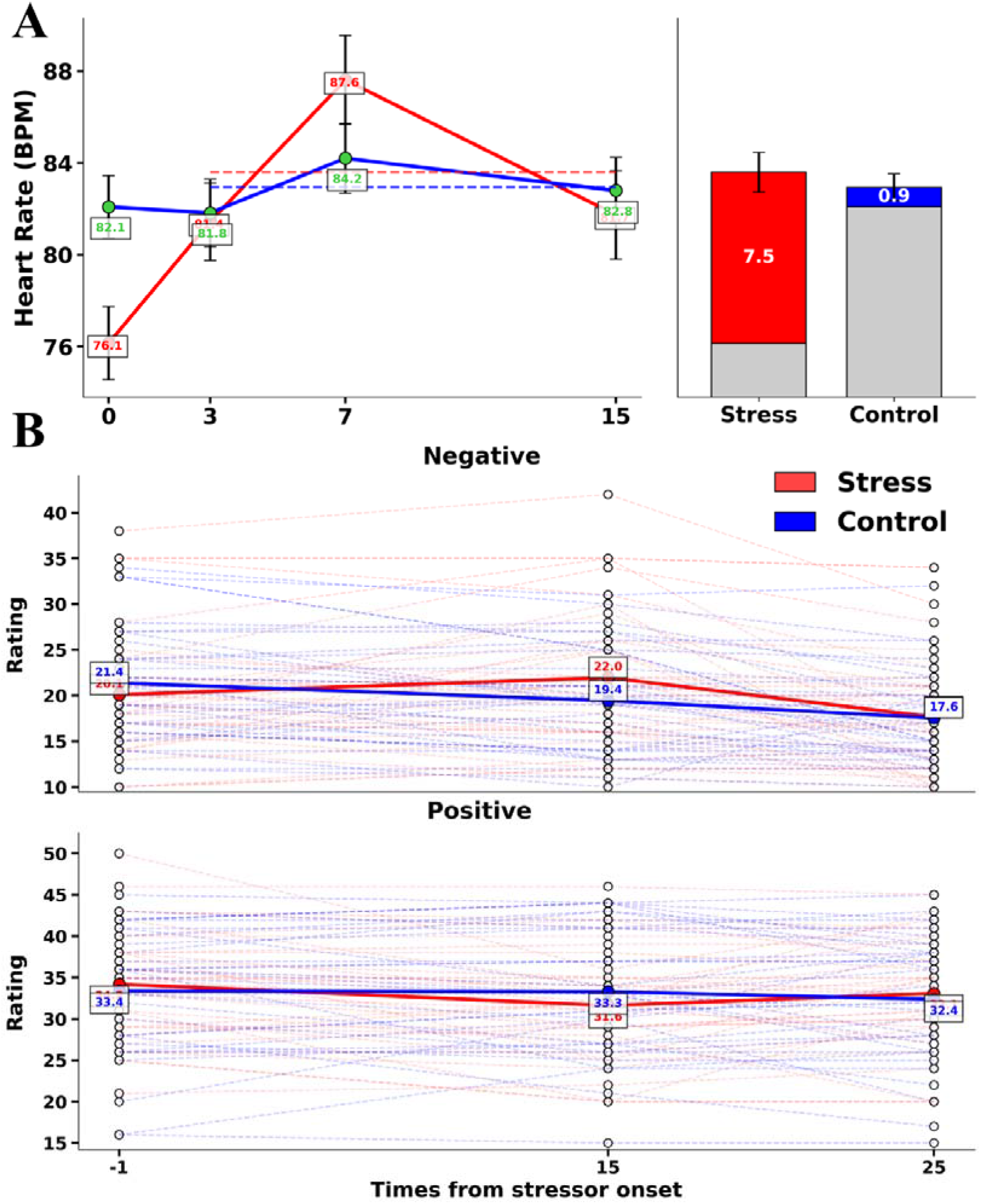
Manipulation Checks. **(A)** Acute stress induction significantly increased heart rate (HR), measured in beats per minute (BPM), during the Trier Social Stress Test (TSST). The heart rate response was defined as the mean BPM difference between the TSST and baseline phases, with the stress group showing a mean increase of 7.5 BPM (SD = 5.27) and the control group showing a mean increase of 0.9 BPM (SD = 3.47). **(B)** Stress significantly increased subjective ratings of negative valence. **(C)** Stress significantly decreased subjective ratings of positive valence.

**Figure S3.**
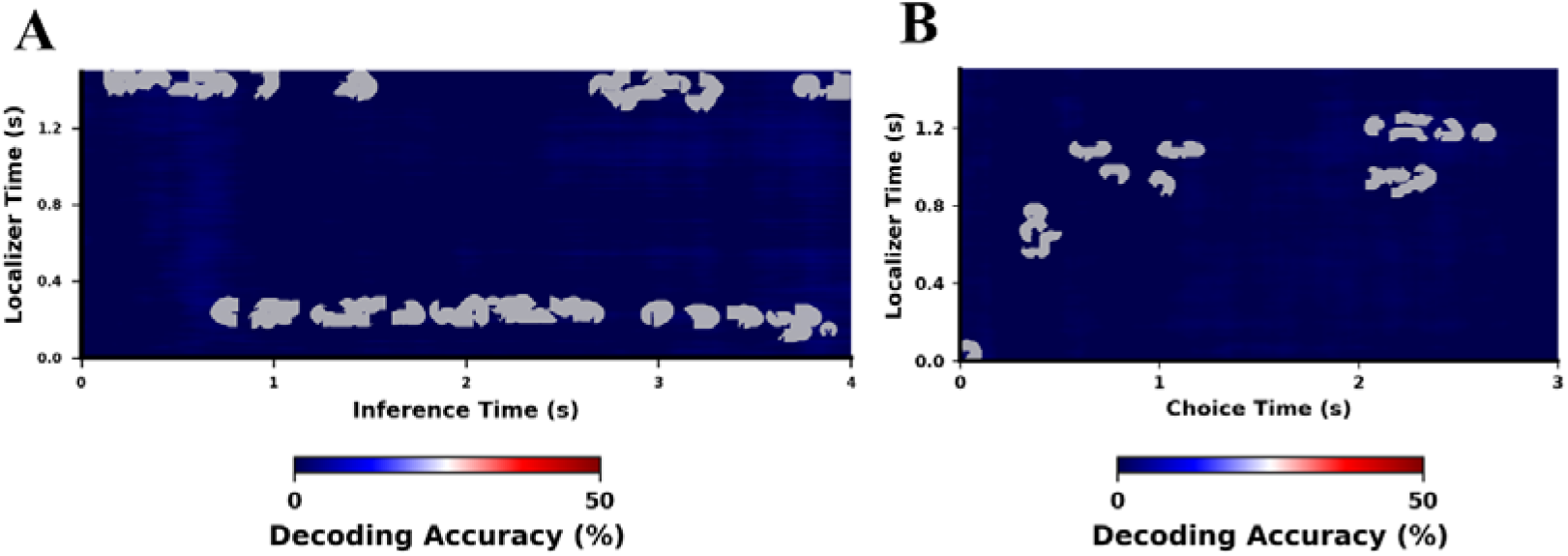
Neural reactivation of bridge-related category information during inference and choice. **(A)** Reactivation during the Inference Phase. The matrix shows generalization to the Inference phase (x-axis), where only the cue (A) was presented. Two primary clusters of reactivation were identified: one extending from early localizer processing (120–290 ms) to a broad inference window (740–3890 ms), and another mapping late localizer processing (1310–1500 ms) to distinct early (90–1500 ms) and late (2680–4000 ms) inference intervals. **(B)** Reactivation during the Choice Phase. The matrix shows the generalization of classifier patterns from the Localizer phase (y-axis) to the Choice phase (x-axis). Significant reactivation of bridge-related category information was observed during the presentation of the cue (A) and options, characterized by early temporal clusters (330–480 ms; 580–1110 ms) and a sustained late cluster (2030–2660 ms).

**Table S1.**
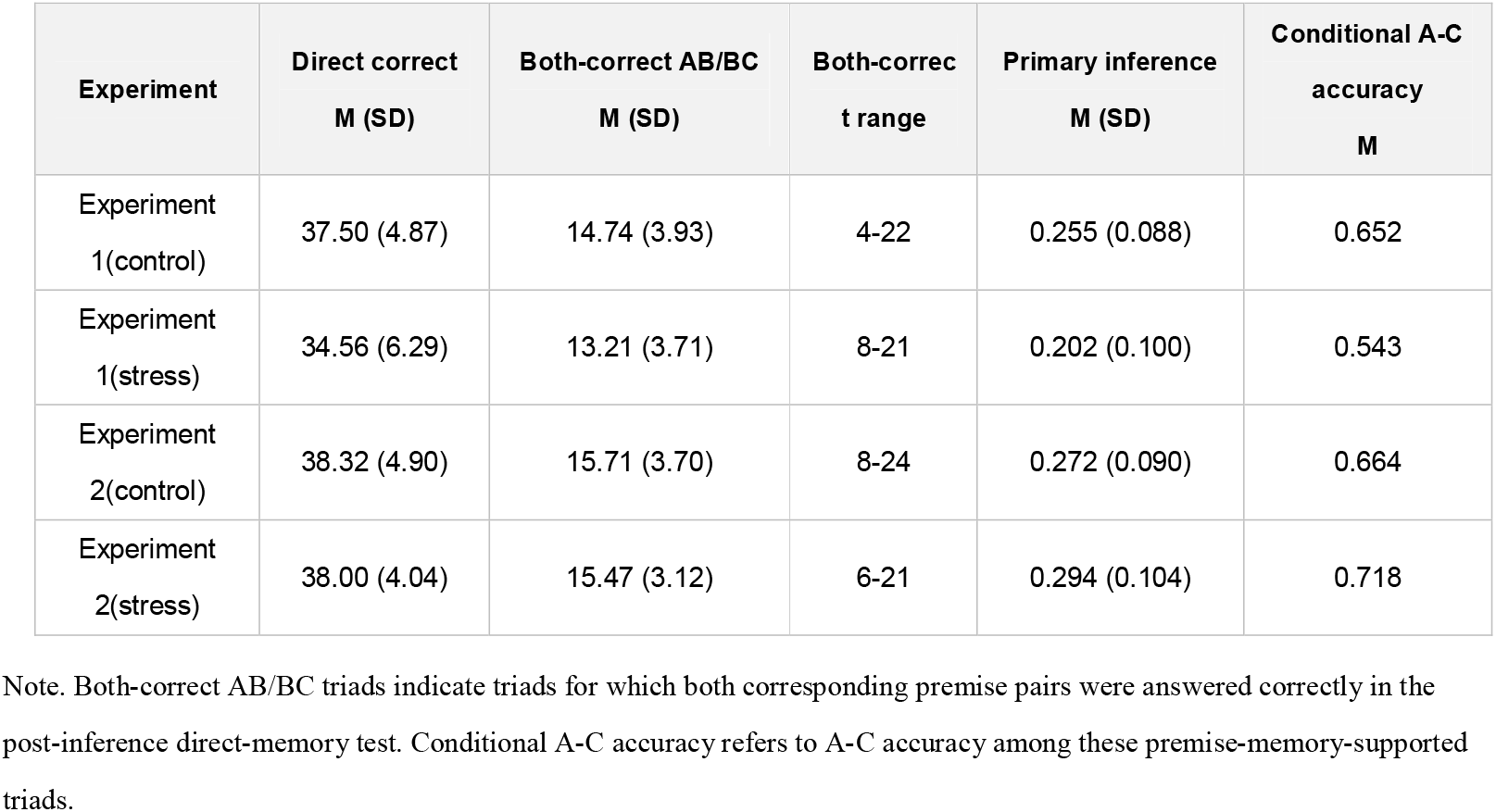
Group-level trial counts.

