## Supplemental Tables and Figures for "Retrieval practice is associated with attenuated stress-related inference impairment and stronger reactivation of bridge-related category information"

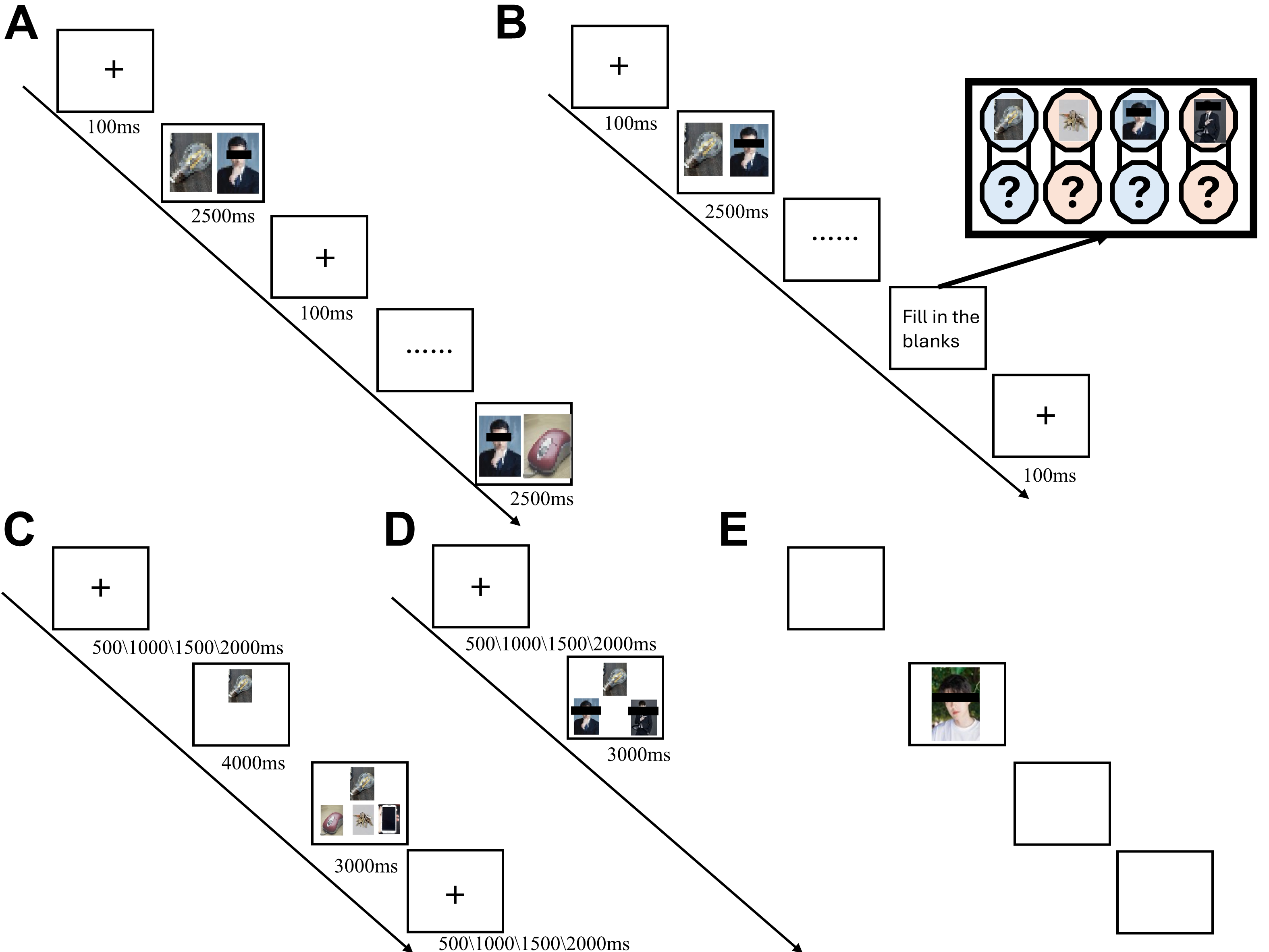

**Figure S1. Experimental procedures.(A) Encoding phase.** Each trial began with a fixation cross (100 ms), followed by the presentation of an AB or BC image pair for 2500 ms. Each pair was presented twice, with the spatial positions reversed in the second presentation. **(B) Retrieval practice phase.** Following the encoding phase, participants were instructed to freely recall and fill in the blanks for the learned memory pairs (AB and BC) using paper and pen within a two-minute time limit. **(C) Memory inference phase.** Each trial started with a fixation point presented for a jittered duration (500, 1000, 1500, or 2000 ms), followed by a memory cue (Item A) for 4000 ms. Subsequently, three options were displayed, and participants were required to select the correct target (Item C) within 3000 ms. Distractors were images from the encoding phase (Day 1) that were not paired with the current cue. **(D) Direct memory retrieval phase.** After a jittered fixation, a cue (Item A or C) and two options (one being the correct Item B) were presented simultaneously. Participants had 3000 ms to make a selection. The distractor was a previously learned image from the same category as the correct option. **(E) Localizer phase.** Images were presented for 1500 ms with a 500 ms inter-trial interval. Participants performed a 1-back task, indicating whether the current image was identical to the preceding one.

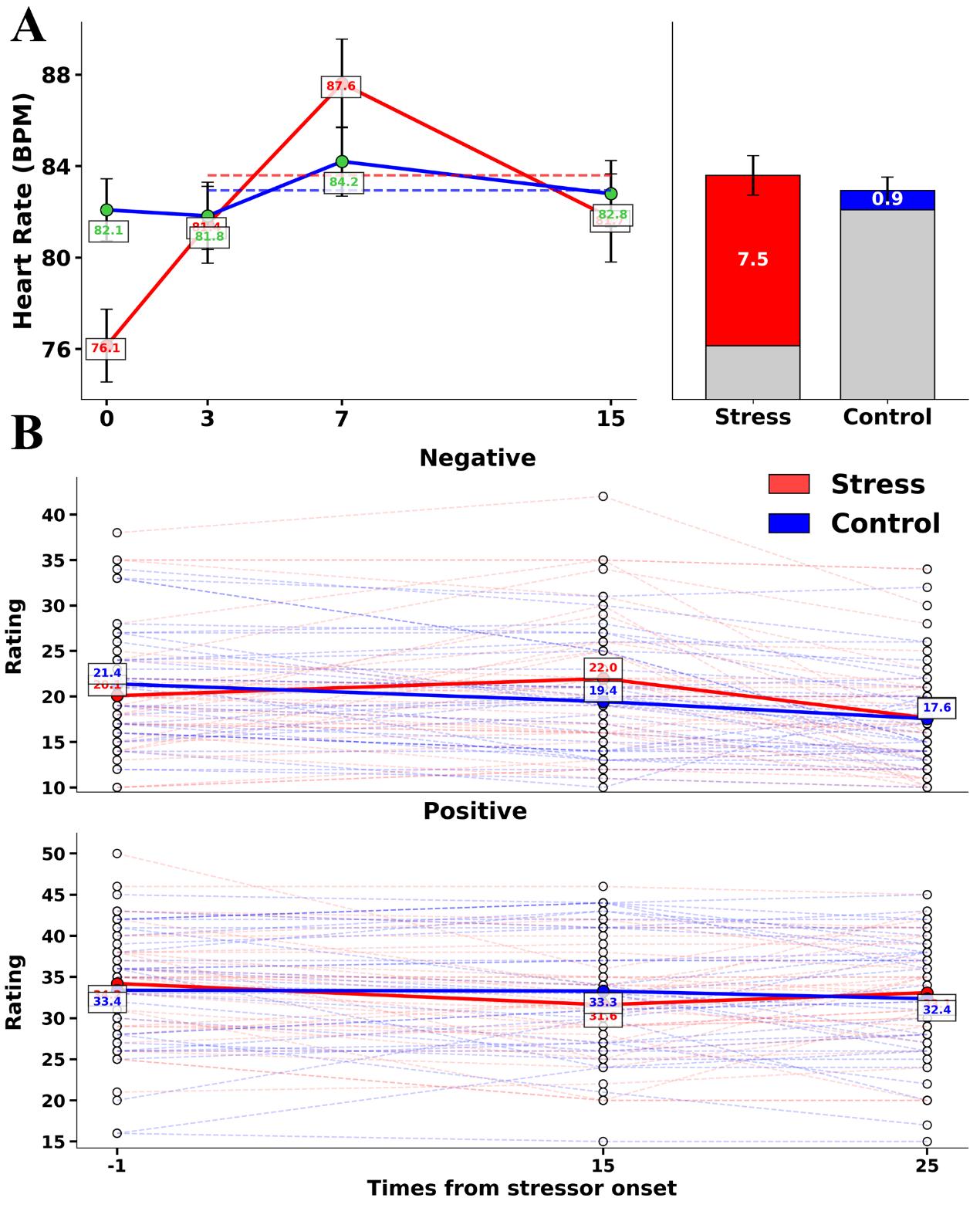

**Figure S2 Manipulation Checks.** **(A)** Acute stress induction significantly increased heart rate (HR), measured in beats per minute (BPM), during the Trier Social Stress Test (TSST). The heart rate response was defined as the mean BPM difference between the TSST and baseline phases, with the stress group showing a mean increase of 7.5 BPM (SD = 5.27) and the control group showing a mean increase of 0.9 BPM (SD = 3.47). **(B)** Stress significantly increased subjective ratings of negative valence. **(C)** Stress significantly decreased subjective ratings of positive valence.

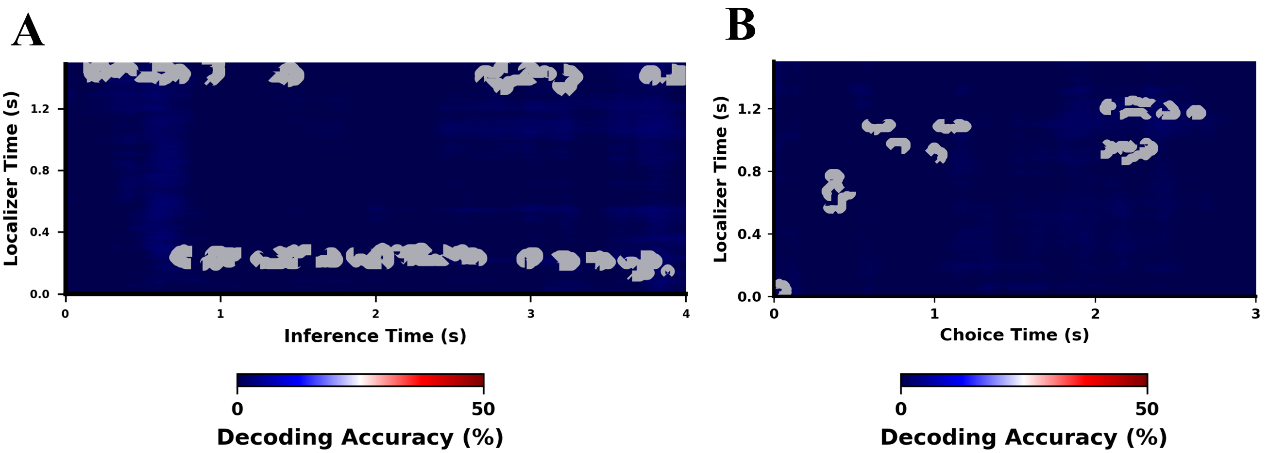

**Figure S3. Neural reactivation for the bridge element (Item B) during inference and choice phases.** **(A)** Reactivation during the Inference Phase. The matrix shows generalization to the Inference phase (x-axis), where only the cue (A) was presented. Two primary clusters of reactivation were identified: one extending from early localizer processing (120–290 ms) to a broad inference window (740–3890 ms), and another mapping late localizer processing (1310–1500 ms) to distinct early (90–1500 ms) and late (2680–4000 ms) inference intervals. **(B)** Reactivation during the Choice Phase. The matrix shows the generalization of classifier patterns from the Localizer phase (y-axis) to the Choice phase (x-axis). Significant reactivation of the bridge element was observed during the presentation of the cue (A) and options, characterized by early temporal clusters (330–480 ms; 580–1110 ms) and a sustained late cluster (2030–2660 ms).

**Table S1. Group-level trial counts.**

| **Experiment** | **Direct correct M (SD)** | **Both-correct AB/BC  M (SD)** | **Both-correct range** | **Primary inference M (SD)** | **Conditional A-C accuracy M** |
| --- | --- | --- | --- | --- | --- |
| Experiment 1(control) | 37.50 (4.87) | 14.74 (3.93) | 4-22 | 0.255 (0.088) | 0.652 |
| Experiment 1(stress) | 34.56 (6.29) | 13.21 (3.71) | 8-21 | 0.202 (0.100) | 0.543 |
| Experiment 2(control) | 38.32 (4.90) | 15.71 (3.70) | 8-24 | 0.272 (0.090) | 0.664 |
| Experiment 2(stress) | 38.00 (4.04) | 15.47 (3.12) | 6-21 | 0.294 (0.104) | 0.718 |

Note. Both-correct AB/BC triads indicate triads for which both corresponding premise pairs were answered correctly in the post-inference direct-memory test. Conditional A-C accuracy refers to A-C accuracy among these premise-memory-supported triads.
